# Exploration of Structural Optic Nerve Changes in Mouse Models of Retinal and Neuronal Degeneration with Optical Coherence Tomography

**DOI:** 10.64898/2026.04.10.717687

**Authors:** Georg Ladurner, Marco Augustin, Danielle J. Harper, Sybren Worm, Maria Varaka, Lucas May, Yash Patel, Theresa Rohrmoser, Fernando García-Ramírez, Gerhard Garhöfer, Manuela Prokesch, Bernhard Baumann, Conrad W. Merkle

## Abstract

**Purpose:** The optic nerve head (ONH) is a central feature of the retina, affected in many human ocular pathologies, yet it has remained underexplored in most mouse models of disease. We hypothesize that the analysis of the ONH can yield valuable insight into the phenotype of retinal diseases and that pathological changes can be detected using state-of-the-art optical coherence tomography (OCT).

**Methods:** Four mouse models – the 5xFAD, PS19 and APP/PS1 models of Alzheimer’s disease (AD) as well as the SOD1 knockout mouse model – were imaged using a polarization-sensitive OCT system to investigate potential disease related changes of the ONH. 5xFAD and SOD1 animals were investigated longitudinally to study disease progression. Additionally, aging effects in wild type mice were studied.

**Results:** Two different analysis methods for the segmentation of the ONH were implemented and evaluated. Longitudinal changes to the ONH in 5xFAD animals were observed, specifically an increase of ONH volume from 3 to 5 months of age followed by a strong decrease until 9 months of age. Significant differences between transgenic (tg) and non-transgenic (ntg) animals, as well as sex dependent distinctions were found. Also, for the APP/PS1 model disease related differences between ntg and tg APP/PS1 were significant.

**Conclusions:** We demonstrated a simple segmentation of the ONH structure based on OCT intensity images and show its potential as a preclinical biomarker in amyloid mouse models of AD.

## Introduction

The optic nerve (ON) and its endpoint in the retina, the optic nerve head (ONH), are crucial structures in the ocular anatomy. The trajectory of the retinal ganglion cell axons leads away from the retinal periphery to the ONH where they can exit the eye through the scleral canal (SC). Through the SC, retinal ganglion cell axons connect to the ON, and retinal and venous blood vessels pass into the eyeball.^1^ The SC thus constitutes the functional bridge facilitating the communication of visual information and nutrient supply.^2^ Pathological changes to the ONH, SC and ON can occur due to a multitude of retinal diseases. Thyroid eye disease alters the blood flow at the ONH.^3^ In glaucoma, optic disk hemorrhage decreases microvasculature, thus changing both ONH anatomy and perfusion.^4, 5^ Additionally, changes in cupping have been reported in relation to irreversible axonal loss.^6^ Reduced vascular density around the ONH occurs in diabetic retinopathy, alongside other vascular disturbances.^7^ Alterations in ONH anatomy have been reported in dry age-related macular degeneration (AMD), where the cup-to-disc ratio and cup shape are subject to pathological changes.^8^ Additionally, reduced ON signal conduction in wet AMD patients has been reported, indicating functional impairment.^9^ The ON and ONH therefore are important targets for the diagnosis of ophthalmic diseases involving retinal degeneration.

In Alzheimer’s disease (AD), degeneration of the ON complements the already well described retinal phenotype of the disease.^10^ As in the brain,^11^ axonal degeneration can be observed in the ON of AD patients in combination with the loss of retinal ganglion cells.^12, 13^ One hypothesis describes the ON as the transport path of amyloid-beta (Aβ) protein from the brain of AD patients to the eye and retina. These findings are supported by histology, confirming the presence of Aβ in the ON of the 5xFAD mouse models and human AD patients.^14^ These findings suggest a potential involvement of the ONH in the retinal pathology of AD, although this type of study has not been performed so far in AD mouse models of any kind.

One important technology for retinal imaging is optical coherence tomography (OCT).^15, 16^ In OCT, the interference-based detection of low coherent light scattered by the sample enables the generation of 3D images of tissue in real time.^17^ State-of-the-art OCT systems have been used to study structural,^18^ topographic,^19^ and vascular ^20^ changes related to disease. Polarization sensitive OCT (PS-OCT) additionally enables the study of pathological changes in optical tissue properties, such as reduced birefringence of the retinal nerve fiber layer in glaucoma ^21^ and diabetic retinopathy,^22^ or changes to melanin containing structures such as the retinal pigment epithelium (RPE).^23^ While OCT has become a routine tool in ophthalmic diagnostics in human patients, OCT studies of the ONH in mice are missing in models of AD, leaving a potentially crucial parameter for preclinical research unexploited.

In this article, we propose two approaches for the structural assessment of ONH geometry in mice and applied them for longitudinal investigations exploring retinal disease progression. Four transgenic mouse models were investigated using a high-resolution PS-OCT system: The 5xFAD, PS19 and APP/PS1 mouse models of AD were used as representative models of neurodegeneration.^24^ Further, the SOD1 mouse model, which can be used to study features of AMD and age-related retinal degeneration,^25^ was investigated to explore the effects of retinal degeneration on the ONH. We hypothesize that the analysis of the ONH can yield valuable insight into the phenotype of retinal diseases and that pathological changes can be detected using state-of-the-art OCT.

## Methods

### Animal Models

#### 5xFAD

The 5xFAD mouse model (The Jackson Laboratory, strain #034848) is a transgenic model of AD, designed to mimic the amyloid pathology found in human AD patients. The model is a common model for preclinical AD studies and has been used in about 10% of all AD mouse studies.^24^ Based on the C57BL/6J background strain, the model is modified with 3 different mutations related to the overexpression of human amyloid-beta precursor protein associated with familial Alzheimer’s diseases (FAD), namely the Swedish (K670N, M671L), Florida 103 (I716V), and London (V717I) mutations. Additionally, the model harbors two FAD mutations to the human presenilin 1 (PS1) protein 104 M146L and 105 L286V. Neurodegeneration appears in the form of cerebral Aβ plaque formation as early as two months of age and loss of synapses and neurons by month 4. Between 8 and 12 months of age, animals experience rapidly increasing plaque number and size in the cortex and hippocampus, coupled with neuroinflammation and stronger neuronal loss.^26^ Male and female 5xFAD animals (32 transgenic (tg), 32 non-transgenic (ntg) littermates) aged 3-9 months and six 22- to 24-month-old non-transgenic 5xFAD animals were provided by Scantox Neuro GmbH (Grambach, Austria).

#### PS19

PS19 mice (The Jackson Laboratory, strain #008169) are a mouse model of tauopathy, overexpressing the human T34 isoform and 4 microtubule binding repeats (1N4R) of the tau protein P301S mutation. These mutations cause tau aggregates followed by cognitive impairment.^27^ Tau aggregates are a major hallmark of AD that have also been observed in the retina of human patients, making this model suited for the investigation of neurodegeneration.^10^ Six PS19 mice (3 tg, 3 ntg) at the age of 39 weeks were provided by Scantox Neuro GmbH (Grambach, Austria).

#### APP/PS1

The APP/PS1 mouse model (Tg(APPswe, PSEN1dE9)85Dbo; The Jackson Laboratory, MMRRC stock number 34829-JAX) is a model of amyloid pathology, expressing a chimeric mouse/human amyloid precursor protein (Mo/HuAPP695swe) and mutant human presenilin (PS1-dE9).^27^ Four tg and 3 ntg animals were acquired from The Jackson Laboratory (Bar Harbor, ME, USA). Animals were investigated at an age of 51-54 weeks.

#### SOD1

The SOD1 knockout (-/-) mouse (The Jackson Laboratory, stock number #002972) lacks the superoxide dismutase (SOD1) system that protects the retina from oxidative damage. The lack of the SOD1 isoenzyme, the isoenzyme with the highest activity in the retina, has been demonstrated to lead to drusen, choroidal neovascularization and retinal pigment epithelium functional loss. ^25^ SOD1 mice have been used as a model of type 3 exudative age-related retinal degeneration, particularly for studies of oxidative stress in AMD. ^28^ SOD1^-/-^ (n=7) with SOD1^+/+^ (n=6) and SOD1^+/-^ (n=3) as control animals were acquired from Jackson Laboratory (Bar Harbor, ME, USA). Details on the study design are displayed in Figure 1.

**Figure 1.**
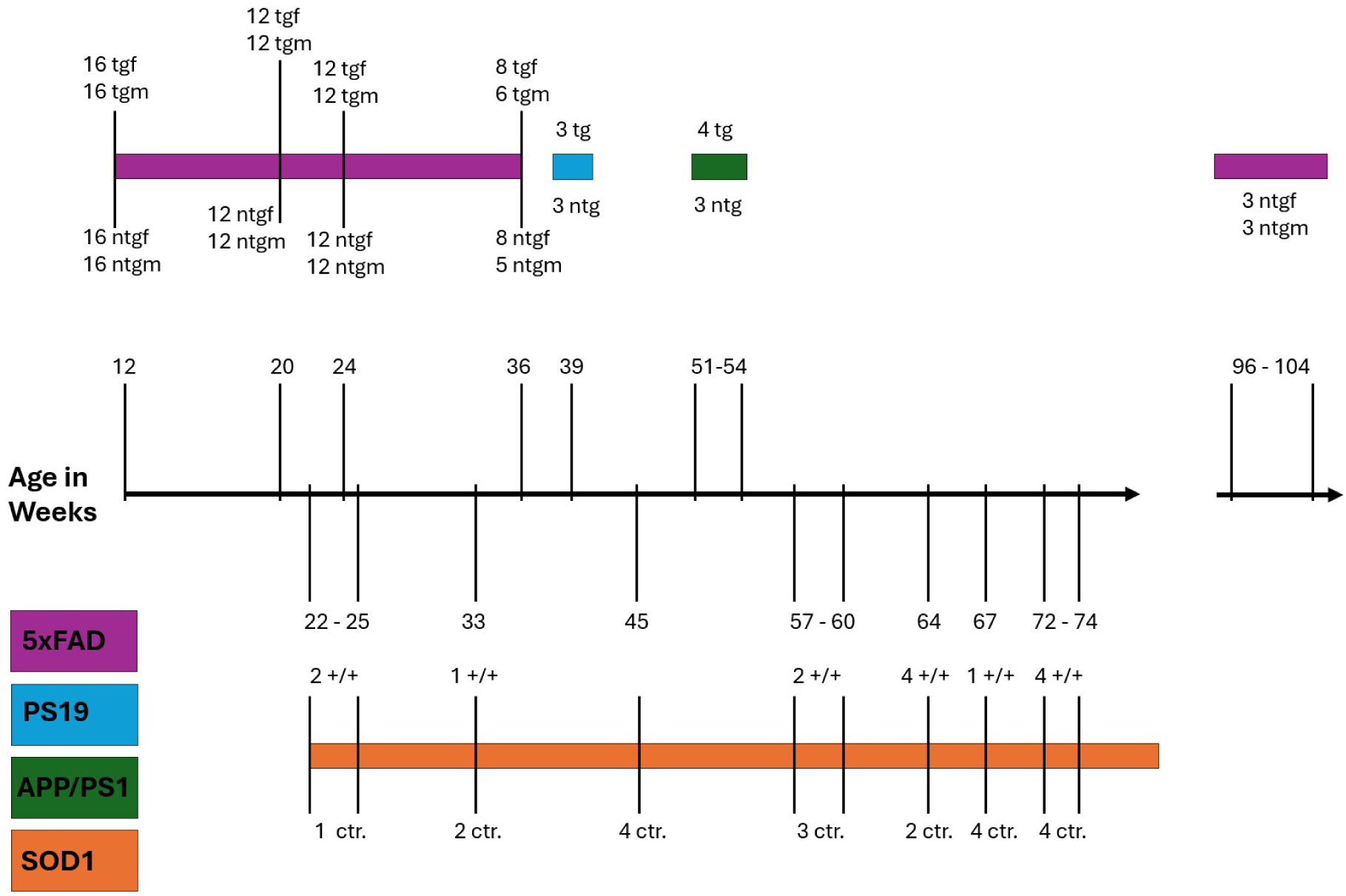
Age and number of animals used for each model with the respective age indicated alongside a time axis. Colors indicate the models as per the legend at the lower left. Transgenic mouse number are printed above the bars, non-transgenic mouse numbers below the bars. Non-transgenic = ntg, transgenic = tg, female = f and male = m, ctr. = control.

Animals were housed in the Core Facility Laboratory Animal Breeding and Husbandry at the Medical University of Vienna under controlled light conditions (12 hours dark, 12 hours light) with food and water ad libitum. Health status and weight was monitored regularly. All experiments were performed in accordance with the ARVO Statement for the Use of Animals in Ophthalmic and Vision Research and Directive 2010/63/EU and with approval by the ethics committee of the Medical University of Vienna and the Austrian Federal Ministry of Education, Science, and Research (GZ: 2024-0.044.300 and GZ: BMBWF-66.009/0216-V/3b/2018).

### Study Design

Two animal models were longitudinally investigated, the 5xFAD and SOD1 knockout mouse. 5xFAD mice were investigated with OCT over the course of 6 months, with measurements at 12, 20, 24 and 36 weeks of age. The timespan between 3 and 6 months of age is the period between early adulthood and the onset of strong neuronal loss in regions of amyloidosis.^27^ From 6 to 9 months of age, a stronger progression of the disease has been observed in the form of increasing plaque formation in the hippocampus and cortex.^24^ The exact number of investigated animals is listed in Supplementary Table 1. Note that the animal numbers decreased over the course of the study because some animals were used for another investigation involving post-mortem histology and were extracted from the study at 12 weeks and 24 weeks of age. ^29^

To investigate the effects of age on the ONH parameters, 6 control animals (3f, 3m) aged 96-104 weeks were additionally investigated. These animals were C57BL/6J ntg-littermate controls of 5xFAD mice but were not part of the longitudinal 5xFAD study. Mouse numbers are displayed in Figure 1 and Supplementary Table 1.

PS19 and APP/PS1 mice were imaged at 39 and 51-54 weeks respectively to investigate the ONH parameters in comparison with the 5xFAD mouse model. Mouse numbers are displayed in Figure 1 and Supplementary Table 2.

SOD1^-/-^ (knockout) (n=7) with SOD1^+/+^ (n=6) and SOD1^+/-^ (n=3) mice serving as controls were investigated with OCT from 22 weeks (5.5 months) to 74 weeks (18.5 months) of age. Scans were acquired at 22-25, 33, 45, 57-60, 64, 67, 72-74 weeks of age. The exact number of mice investigated per timepoint is listed in Figure 1 and Supplementary Table 3.

### OCT System

The PS-OCT system presented by Fialová et al. was used for imaging.^30^ The system was powered by a superluminescent diode with a central wavelength of 840 nm and a spectral bandwidth of 100 nm. The axial resolution amounted to ∼3.8 µm in tissue (n=1.35). Spectral data was acquired using spectrometer line scan cameras acquiring 3072 pixels for the co- and cross-polarized channel, operating at an A-scan rate of 80 kHz for 5xFAD and PS19 mice and 83 kHz for APP/PS1 and SOD1 mice. A stage with three translational and two rotational degrees of freedom was used to align the mouse eye, so that the field of view, measuring around 1×1mm, was centered on the ONH. Volumes composed of 2000 B-scans, each consisting of 512 A-lines, were acquired by imaging five repeated B-scans at 400 slow axis positions. The resulting volumes used for analysis consisted of 5×512×400×3072 spectral voxels.

### Anaesthesia and Imaging

The imaging protocol applied in this study followed the recommendations made by Ladurner et al., 2025.^31^ Experiments were performed during the 12-hour daytime period of the animals. All mouse models were imaged under isoflurane anaesthesia. Mice were placed in an induction chamber that was subsequently flooded with 4% isoflurane (IsoFlo, Zoetis Österreich GmbH) in oxygen for 4 minutes. Next, the mice were moved to the home-built imaging stage, where 2% isoflurane anaesthesia was supplied through a 3D printed nose cone. Some animals were more resistant to isoflurane after repeated anaesthesia requiring a concentration increase to 2.5%. Mouse eye lid reactions were monitored throughout the experiment to ensure proper depth of anesthesia. Tropicamide drops (0.5%, Agepha Pharma s.r.o., Senec, Slovakia) were applied to dilate the pupils for imaging. A heating blanket was used to control mouse body temperature and prevent hypothermia. Oculotect eye drops (Théa Pharma GmbH, Berlin, Germany) were repeatedly applied to hydrate mouse eyes during the entire imaging session. The drops were removed using cotton swabs prior to image acquisition to eliminate lensing effects induced by the fluid film on the eyes. Both eyes of each animal were imaged without interrupting the anaesthesia. Time under anaesthesia was kept below 40 minutes per imaging session. For most animals, the time between anaesthesia induction and end of the imaging session was between 12 and 35 minutes. A heating pad was provided to support the mouse during anaesthesia recovery.

### Image Processing

Basic OCT image processing was performed using the pipeline previously described.^32^ The polarization contrast provided by the system was used to detect the boundaries of the retinal pigment epithelium (RPE) via the strong depolarizing properties of the melanin pigments contained in this layer.^33, 34^ A previously presented algorithm based on edge detection was hereby used to segment the RPE in the cross-polarized channel.^32, 35^ The curvature of the retina was then flattened based on the axial positions obtained from the RPE segmentation, and interframe motion was removed by aligning the frames with respect to the flattened RPE. The data analyzed for 5xFAD mice,^29^ PS19 and APP/PS1 mice,^31^ as well as the SOD1 mice^36^ had been acquired and evaluated in previous studies. Reusing data helps to reduce the number of animals used in studies overall and aids the implementation of the 3Rs of animal research (Replacement, Reduction, and Refinement). Here all data sets were re-processed with the updated and advanced pipeline described below to investigate and quantify the ONH structure for all mouse models. Data sets that could not be processed due to strong vignetting, low signal quality, partial obstruction of the field of view by cataract, acquisition errors or others were excluded from the study.

In the following, we will present two methods for the quantification of ONH structure. One is based on ONH cross-sectional areas and applied for different depths of the OCT B-scan. In the second approach we compose volumes from segmented areas within a given depth range.

### Analysis of Cross-Sectional ONH Area with Adapted Threshold

To segment cross-sectional areas, we chose the anterior and posterior boundary of the RPE as our reference planes, as the RPE boundaries in the OCT-volume show elevated contrast compared to other layers. As the ONH region is generally devoid of contrast at the level of the outer retina, the segmentation in the ONH region was thus determined by the pixels of the surrounding RPE boundary. This guaranteed that the analysis was always performed in the same ONH region. Figure 2A shows the segmentation applied on one B-scan. The segmentation of retinal layers was performed according to Augustin et al., 2018.^35^ The coordinates from the segmentation of the anterior and posterior RPE were mapped to the corresponding pixels in log scale intensity volumes in order to extract 2D reflectivity maps for the anterior and posterior RPE, respectively. Each of these reflectivity maps were slices that extended over one pixel in depth. The resulting slices in Figure 2C show the intensity distribution depending on the chosen depth. In Figure 2, slices at the anterior and posterior RPE boundary are shown, as well as planes 10 and 20 pixels (20 and 40 µm) below the posterior RPE boundary. The respective positions of these four planes are visualized in a B-scan in Figure 2B. To extract the ONH cross-sectional area from the intensity maps, first median filtering using a 5×5 pixel kernel for smoothing and then intensity thresholding was applied. In each map, all pixels with a threshold in log scale intensity (TI) 2.5 times the standard deviation (STD) below the mean (𝑇𝐼_2.5_ = 𝑀𝑒𝑎𝑛(𝐼) − 2.5 ∗ 𝑆𝑇𝐷(𝐼)) were classified as ONH. For the plane 40 µm below the posterior RPE, 𝑇𝐼_2.5_ was not optimal, hence TI was optimized to 2.0 times the STD below the mean (𝑇𝐼_2.0_ = 𝑀𝑒𝑎𝑛(𝐼) − 2.0 ∗ 𝑆𝑇𝐷(𝐼)). Figure 2D shows the pixels classified as ONH in grey, superimposed over the false-color intensity planes. ONH area measurements were then obtained by summing the segmented pixel count and scaling by the effective pixel size of 5 µm².

**Figure 2.**
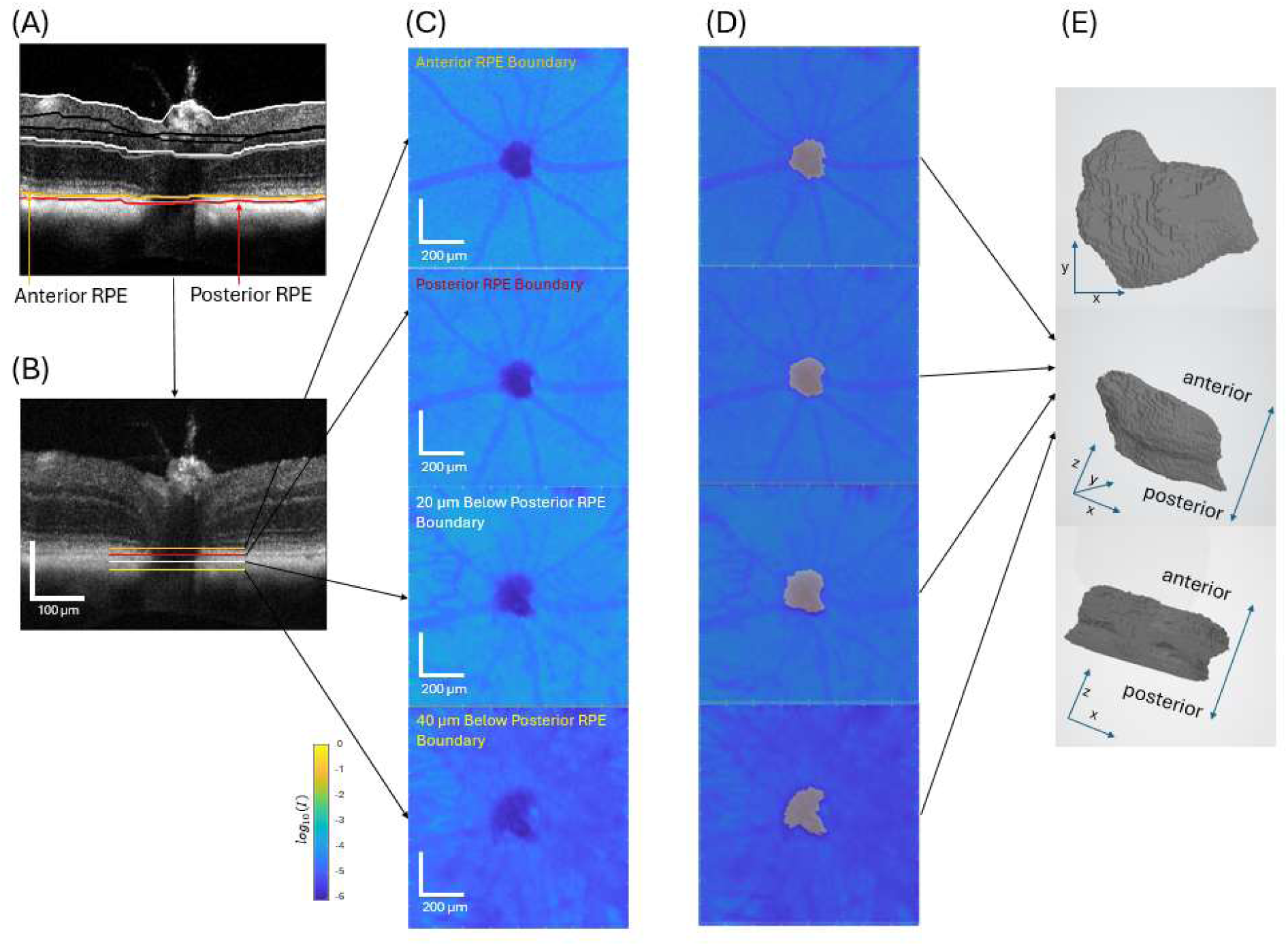
Segmentation of retinal layers and ONH. (A) Layer segmentation of anterior and posterior RPE boundary shown on one B-scan. This segmentation was used for the depth positioning. (B) Representative position of depths used for ONH volume segmentation in a flattened B-scan: anterior and posterior RPE boundary, as well as 20 and 40 µm below the posterior RPE boundary. (C) En face reflectivity maps in the selected depths positions. (D) Segmented ONH areas in each depth are indicated in grey. (E) Volume rendering of the segmented ONH section.

### Quantification of ONH Volume in a 26-pixel range (52 µm)

To assess the three-dimensional structure of the ONH, we quantitatively analyzed volumes composed of segmented areas stacked on each other. To this end, the posterior RPE boundary was chosen as a reference and all areas within a 52-µm deep range extending from 30 µm above to 20 µm below the posterior RPE were segmented using the method described for areas with 𝑇𝐼_2.5_ as a threshold. Areas were then converted into volume sheets by multiplication with the voxel volume of 10 µm^3^.The limits of this axial range were empirically chosen because this region displayed the highest signal-to-noise ratio and was thus most reliable for the segmentation. Averaged volumes with a larger axial range (142 µm) were also segmented. This data is presented in the Supplement.

### Data Screening

The ONH area segmentation was dependent on signal quality and was thus affected by eye motion or locally low signal. For both segmentation approaches, areas and volumes were therefore manually screened and excluded from further analysis when the segmentations were unreliable. Examples for excluded volume scans with the corresponding intensity maps at the anterior RPE as well as volumes are shown in Supplementary Figure 1. A table with the percentages of removed scans for ONH volumes, per mouse model and imaging timepoint is shown in Supplementary Table 10 and 11.

### Statistical Analysis

Longitudinal data was evaluated using mixed effects models to take the measurement of both eyes of the animals into account. Pairwise comparisons between groups were calculated using estimated marginal means (EMM). Since the sample size and distribution for SOD1 mice was too low for a stable mixed effects model, an ANOVA was used. Pairwise comparisons were extracted using Tukey’s test. The Benjamini-Hochberg correction was used for tg/ntg comparisons. For sex-based analysis, no post correction was performed due to the exploratory character of the evaluation. P-values were considered significant for p<0.05.

## Results

### Analysis of Cross-sectional ONH Areas in 5xFAD Mice

The longitudinal analysis of cross-sectional ONH areas revealed consistent characteristics across all investigated depths, with areas enlarging from 3 to 5 months of age and shrinking from 5 to 6 and 9 months of age. Figure 3 presents the longitudinal development of the ONH cross-section in different depths for the pool of male and female 5xFAD animals. Most notably, compared to ntg litter mates, tg animals exhibited significantly larger ONH areas at the latest timepoint for all depths except 20 µm below the posterior RPE (see p-values in Figure 3). Means and standard deviations for all timepoints can be found in Supplementary Table 5. For the depth 40 µm below the RPE, larger areas were observed in tg animals for all investigated timepoints except for 6 months of age (p=0.256).

**Figure 3.**
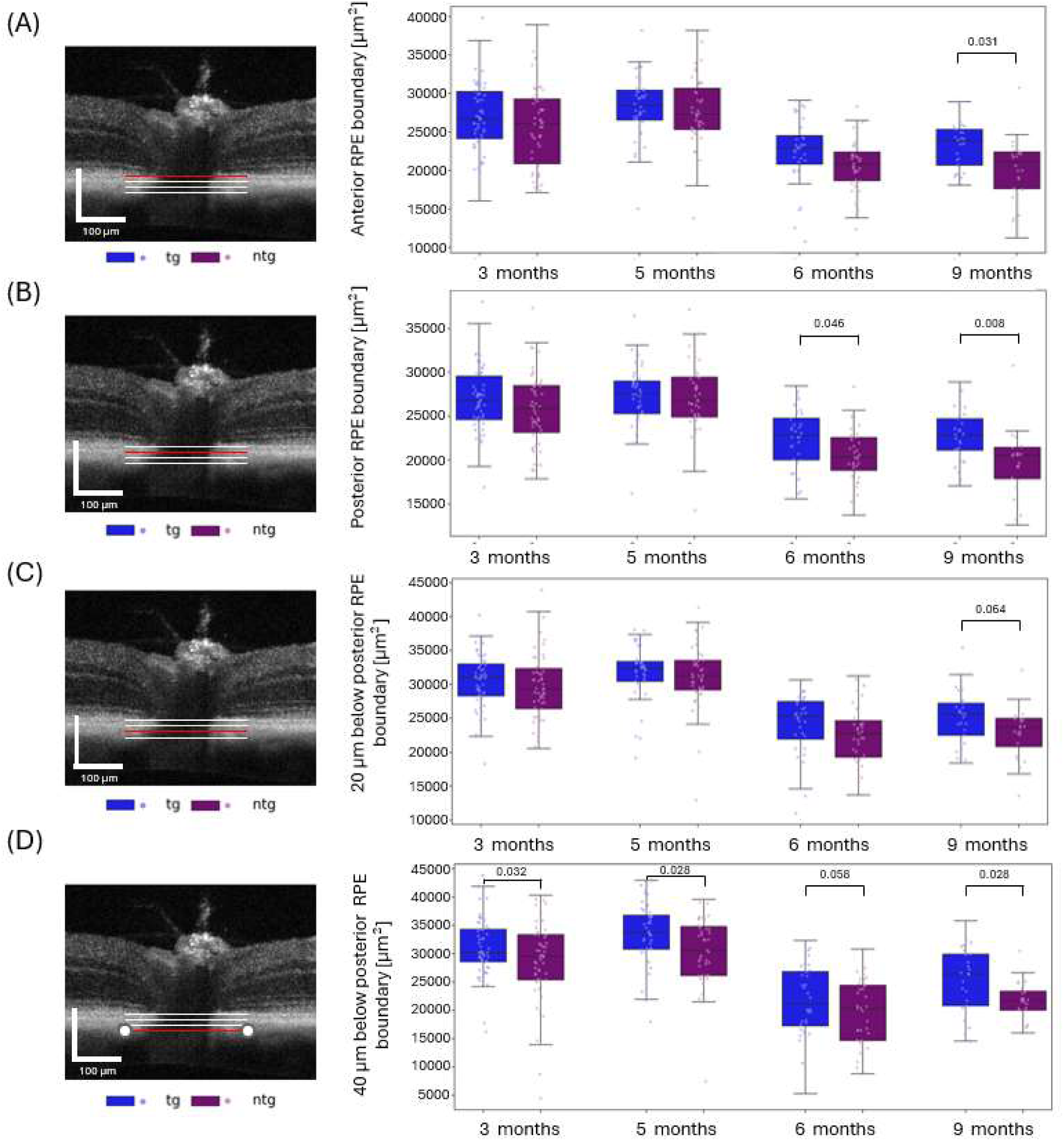
Longitudinal development of ONH cross-sectional area in four different depths for transgenic (tg) and non-transgenic (ntg) 5xFAD animals (male and female subjects pooled): (A) Anterior RPE boundary, (B) posterior RPE boundary, (C) 20 µm below the posterior RPE boundary and (D) 40 µm below the posterior RPE boundary. Boxplots show the median (line), 25-75 percentile (boxes), interquartile range (whiskers), as well as all individual measurements. P-values are indicated above brackets for significantly different data pairs only.

In Figure 4, we present a sex-specific analysis investigating male and female animals separately. Here, at the depth of the anterior RPE, an average non-significant area increase of about 6% was observed between the first measurement (3 months of age) and the second measurement at 5 months of age (Figure 4A). With an overall drop of about 27%, the measured area significantly decreased from 5 to 9 months of age for all mice according to the mixed effects model (p<0.0001). For 9 months of age, the ONH area of tg female (tgf) animals (20204 µm^2^ ± 6763 µm^2^) was 2.6% larger than for non-transgenic female (ntgf) animals (19692 µm^2^ ± 5143 µm^2^) (p=0.011). Significant differences were also observed between tg male (tgm) mice and ntg male (ntgm) mice at 6 months of age. Here the tg animals had 14.2% larger areas with 23628 µm^2^ ± 3470 µm^2^ compared to ntg animals with 20688 µm^2^ ± 4715 µm^2^ (p=0.036). ONH areas also differed significantly between tgm and tgf mice. Similar trends were observed for the ONH area at the depth of the posterior RPE boundary (Figure 4 (B)). Areas increased for all animals from the first to the second timepoint by about 3.3% and subsequently decreased by about 21.5% from 5 to 9 months of age (p=0.067). Significant differences were observed between ntg and tg female animals at 9 months of age, with tgf animals having 26.0% larger areas (24493 µm^2^ ± 11244 µm^2^) than ntgf mice (19445 µm^2^ ± 4723 µm^2^) (p=0.005), as well as for tgm and ntgm animals at 6 months of age, where tgm mice (23458 µm^2^ ± 2838 µm^2^) display 15.1% larger areas than ntgm (20454 µm^2^ ± 3422 µm^2^) animals (p=0.036).

**Figure 4.**
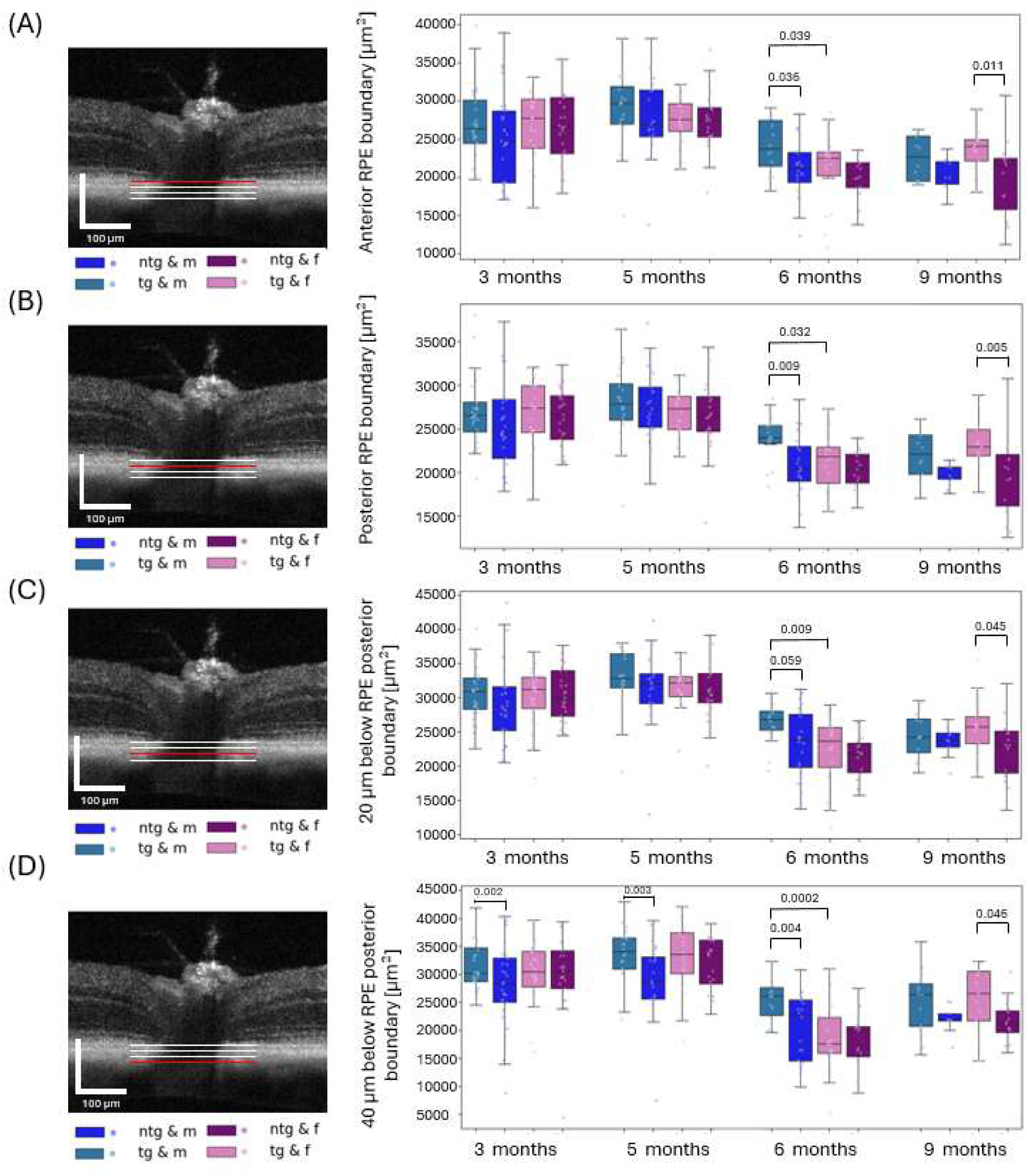
Longitudinal development of ONH areas in four different depths for transgenic female (tgf), transgenic male (tgm), non-transgenic female (ntgf) and non-transgenic male (ntgm) 5xFAD animals: (A) Anterior RPE boundary, (B) posterior RPE boundary, (C) 20 µm below the posterior RPE boundary and (D) 40 µm below the posterior RPE boundary. Boxplots show the median (line), 25-75 percentile (boxes), interquartile range (whiskers), as well as all individual measurements (data points).

ONH areas extracted 20 and 40 µm below the posterior RPE boundary showed a similar behavior. The cross-sections increased from 3 to 5 months of age and then decreased for the later timepoints. Significantly larger areas were found for tgf in comparison to ntgf animals at 9 months of age in both cases (Figure 4(D)). All means and standard deviations can be found in Supplementary Table 1. The analysis of ONH areas assessed 20 µm above the anterior RPE revealed a different behavior compared to the other levels presented above and is shown in Supplementary Figure 2.

### Analysis of ONH Volumes in 5xFAD Mice

Volume calculated from a 52 µm depth range with the highest segmentation reliability were investigated, stratified by genotype and by pooling the data from all tg and ntg 5xFAD animals (Figure 5B and 5C). All volumes increased from 3 to 5 months by about 11 % and decreased significantly by about 21% from 5 to 6 months of age (p=0.005) and by another 9.5% from 6 to 9 months of age (p<0.0001). ONH volumes of tg animals (613874 µm^3^ ± 148670 µm^3^) were on average about 27% larger than for ntg animals (450151 µm^3^ ± 118555 µm^3^) (p=0.0008), as shown in Figure 5C. Of note, male and female mice contributed differently to this effect. With about 25% difference, ONH volumes of tgm animals were substantially but not significantly (p=0.098) larger than those of ntgm mice. For tgf mice, volumes were 32% larger than for ntgf mice (p=0.0004). The volume data from female mice thus contributed more strongly to the observed significant difference between all tg and ntg animals at 9 months of age. As the number of female animals was slightly higher this might also influence the statistics.

**Figure 5.**
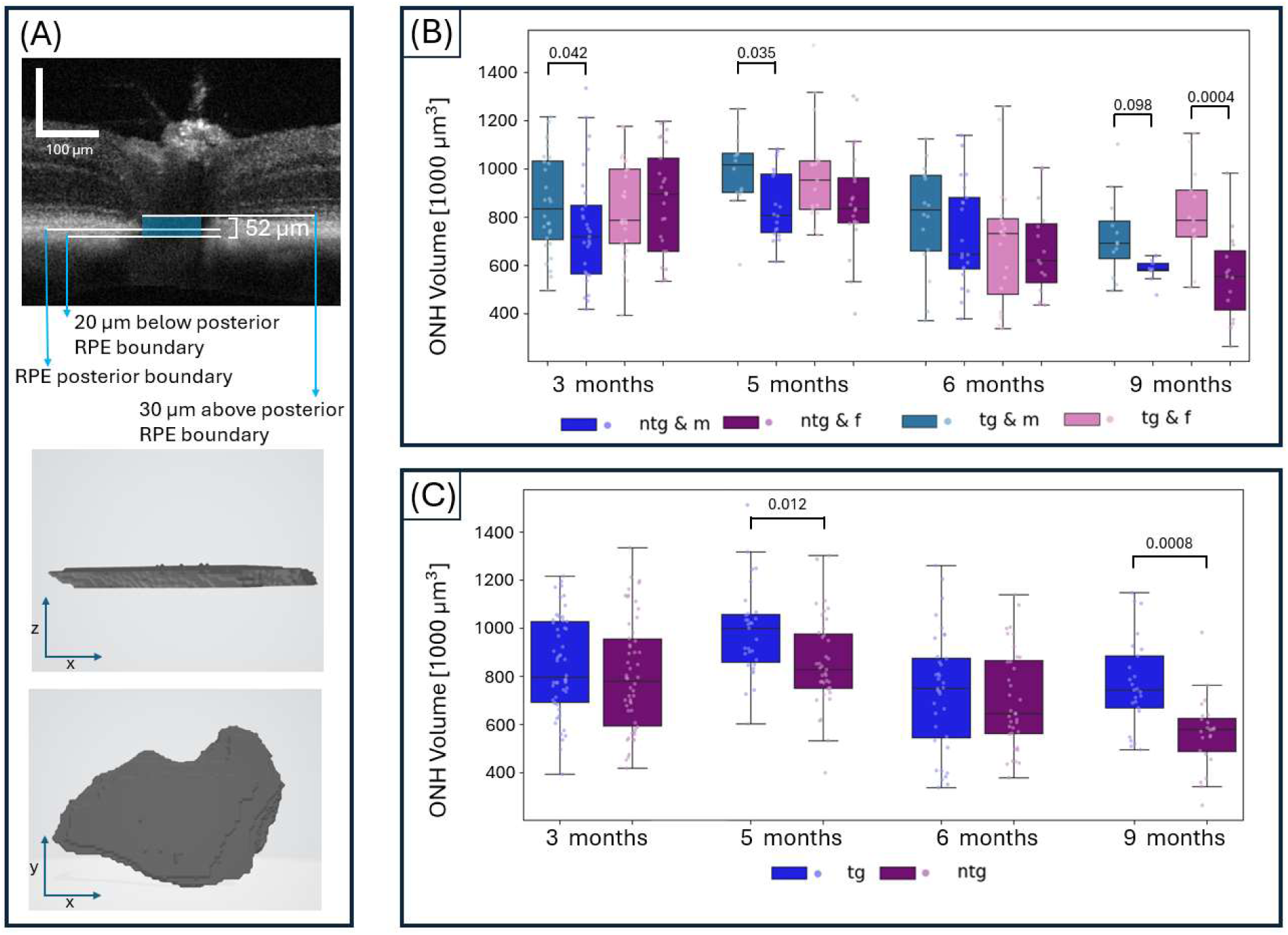
Longitudinal development of ONH volume for 5xFAD mice evaluated in a 52 µm range around the posterior RPE boundary. (A) Visualization of the evaluated depth range in a B-scan and exemplary rendering of the segmented structure. (B) Longitudinal ONH volume stratified for sex and genotype. (C) Longitudinal ONH volume for ntg and tg animals after pooling female and male subjects. Boxplots show the median (line), 25-75 percentile (boxes), interquartile range (whiskers), as well as all individual measurements.

While tgm and ntgm animals also presented with significant volume differences for 3 and 5 months of age, in female mice significant differences were observed exclusively at 9 months of age. For the pooled data of all tg and ntg animals, also the comparison at 5 months of age revealed about 13% larger volumes for tg animals (p=0.012).

Volumes measured for an extended depth range of 142 µm, are described in the Supplement with Supplementary Figure 3 showing the sex stratified and pooled results for the 5xFAD animals. Means and standard deviations for shown values are listed in Supplementary Tables 4 and 5. The behavior observed for small ONH volumes (52 µm) was similarly present for extended volumes (142 µm), with a volume increase from 3 to 5 months of age, followed by a steady decrease until 9 months of age. Significant differences between tg and ntg animals were found for 6 and 9 months of age (p=0.044 and p=0.012).

### Long Term Aging in Control Animals

To investigate how the ONH volume (26 pixel/52µm range) develops from early adulthood to very old age, an additional group of 6 ntg animals was imaged at the age of 22-24 months, thereby complementing the already presented 5xFAD data. Figure 6 shows the investigated region (Figure 6A) and the longitudinal development of the ONH volume for female and male mice (Figure 6B) as well as for all ntg animals (Figure 6C). At 3 months of age, ONH volumes were significantly larger in male animals than female mice (p=0.033). For all other timepoints, no significant differences were observed, indicating that this parameter was not dependent on sex in control mice.

**Figure 6.**
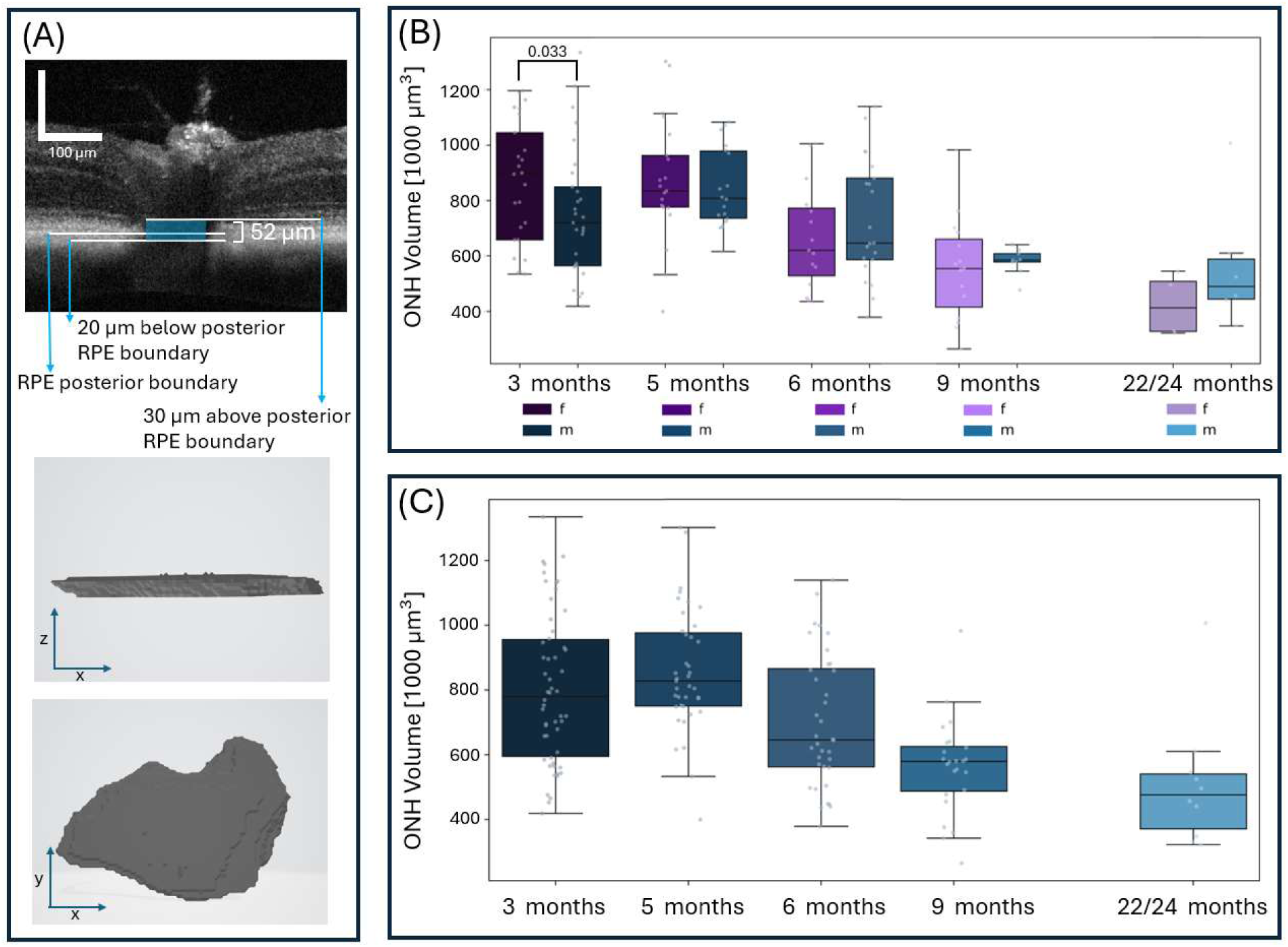
Longitudinal development of ONH volume for ntg control mice evaluated in a 52-micron range around the posterior RPE boundary (A) for female and male animals (B) as well as all animals (C). Boxplots show the median (line), 25-75 percentile (boxes), interquartile range (whiskers), as well as all individual measurements.

The ONH volume first increased by 6.7% from 3 to 5 months of age from 804,085 µm^3^ ± 233,341 µm^3^ to 858,562 µm^3^ ± 187,195 µm^3^. Then, the ONH volume decreased to 717,432 µm^3^ ± 209,958 µm^3^ at 6 months of age and finally to 562,689 µm^3^ ± 148,193 µm^3^ at 9 months of age. These statistically significant changes amounted to about 16.5% (p=0.046) and 21.6% (p<0.001) reductions from 5 to 6 and 6 to 9 months, respectively. Overall, the average ONH volume shrunk from 5 to 9 months of age by almost 35%. From 9 to 22-24 months of age, the volume again significantly reduced by about 10% to 508,116 µm^3^ ± 199,955 µm^3^ (p=0.030). From 3 months to about 2 years of age, the investigated mice thus lost about 37% of the initial ONH volume and about 41% of the maximum volume at 5 months of age. Please see Supplementary Table 6 for all measured mean values and standard deviations.

### Comparison of ONH Structures in three AD Mouse Models

To explore ONH parameters in other AD mouse models, we also compared the results obtained for the 5xFAD mice with PS19 and APP/PS1 of similar age (36-54 weeks). Figure 7 shows the comparison of the models for the segmented volume (Figure 7A) and the areas at the depth of the anterior and posterior RPE boundary (Figure 7B-C). Observed ONH areas and volumes differed markedly for the two amyloid models and the PS19 mice. 5xFAD as well as APP/PS1 mice showed higher values for tg animals, but not for PS19 animals. The ONH areas differed significantly between tg and ntg animals for both APP/PS1 (p=0.0115 and p=0.0009 for anterior and posterior RPE level) and 5xFAD mice (p=0.0045 and p=0.0011 for anterior and posterior RPE). For 5xFAD animals, also the ONH volumes differed significantly between tg and ntg animals (p=9.39×10^-6^). For PS19 mice, however, the ONH areas and volumes did not differ significantly between tg and ntg subjects. Overall, measured areas and volumes were at least 20% larger for APP/PS1 mice than for 5xFAD or PS19 mice (p-value between 6.66×10^-15^ and 2×10^-6^), while they did not differ significantly in PS19 and 5xFAD. Again, the observed behavior is similar for all methods of ONH segmentation. Means and standard deviations are displayed in Supplementary Table 4. P-values for all comparisons are listed in Supplementary Table 8 and are visualized in Figure 8.

**Figure 7.**
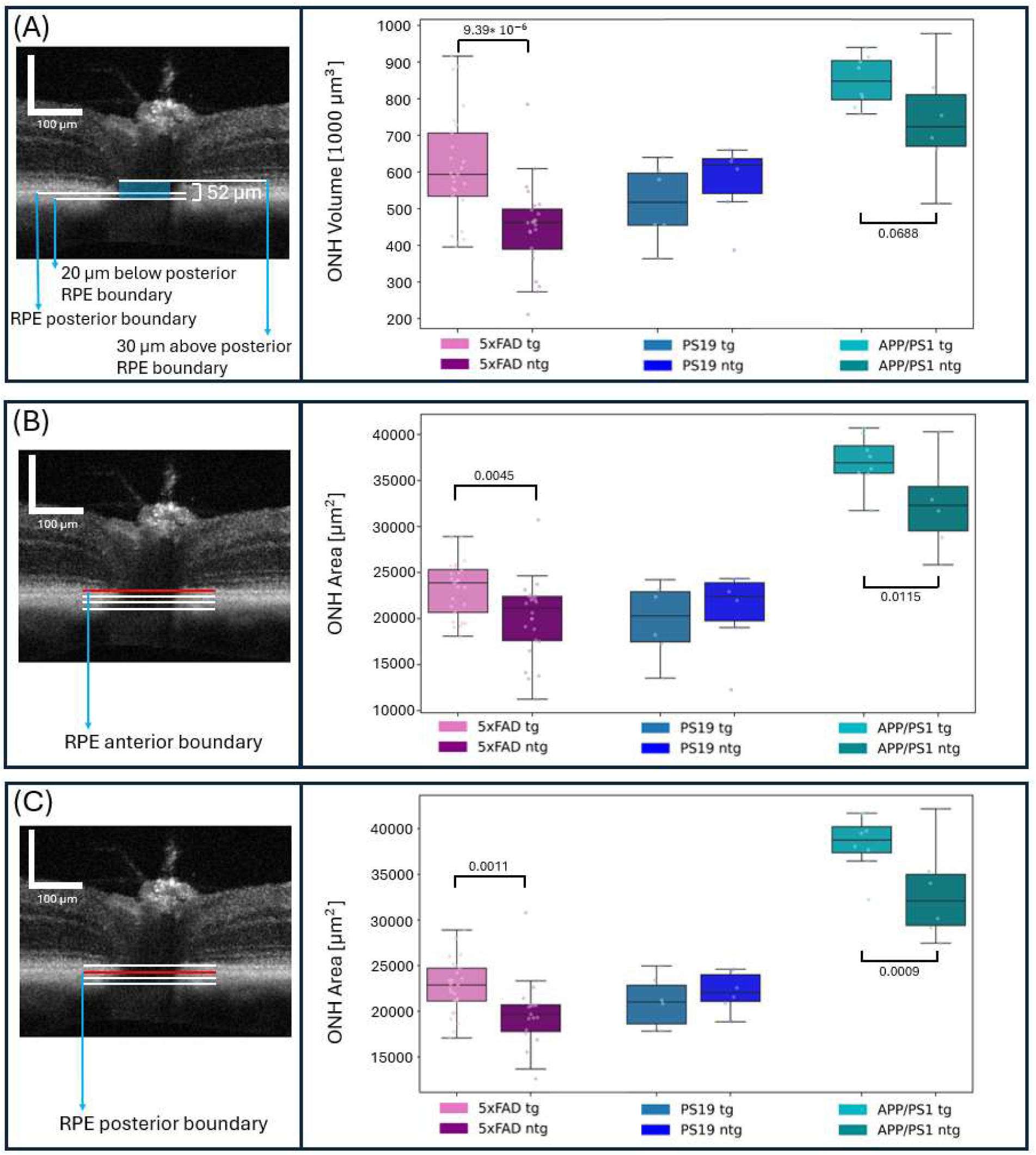
Comparison of ONH volume for tg and ntg mice between 5xFAD, PS19 and APP/PS1 animals (36-54 weeks old). Volume in a 52-micron range around the lower RPE (A) observed area at the depth of the anterior RPE boundary (B) as well as the observed area at the depth of the posterior RPE boundary (C). Boxplots show the median (line), 25-75 percentile (boxes), interquartile range (whiskers), as well as all individual measurements.

**Figure 8.**
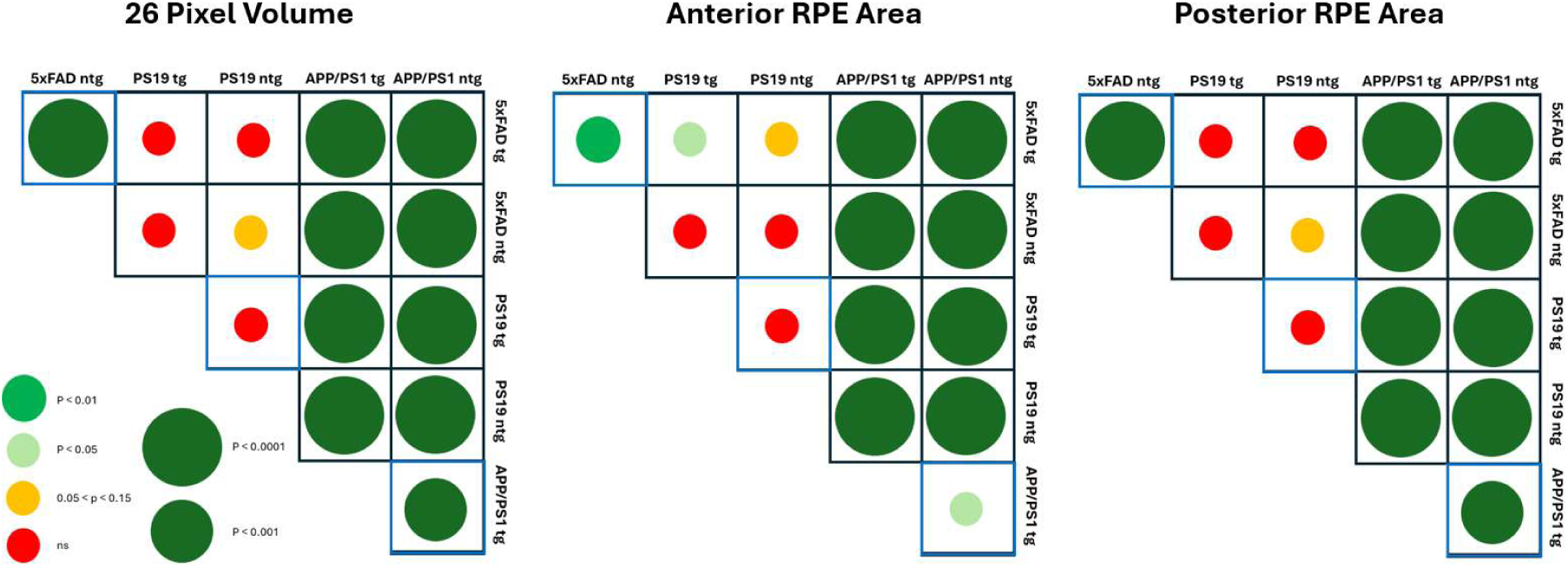
P-values for all comparisons tested between the AD mouse models (aged 36-54 weeks) for the volumes and areas at anterior and posterior RPE depth. Red disk: non-significant (p>0.15), yellow disk: non-significant (0.15>p>0.05), green disks: significant (p<0.05). Disk sizes scale inversely with the p-value as indicated in the legend at the lower left.

### SOD1 Mouse Model

Results from the longitudinal investigation of ONH areas and volumes in the SOD1 mouse model are displayed in Figure 9. In this model, no significant differences between knockout and control animals were observed for the mouse cohorts. A slight yet not statistically significant aging effect was visible for the knockout mice in the form of a shrinking of ONH parameters from 33 weeks onwards. An increase of 15%, 5.6 % and 9.4% was observed for volumes, posterior RPE area and anterior RPE area, respectively, from 22-25 to 33 weeks of age. For measurements at the anterior (Figure 9B) and posterior (Figure 9C) RPE boundary, a loss of area over time appeared more prominent than for the volume analysis. For control animals, an aging effect was not observable in the presented data. Means and standard deviations of the data shown in Figure 9 are listed in Supplementary Table 9.

**Figure 9.**
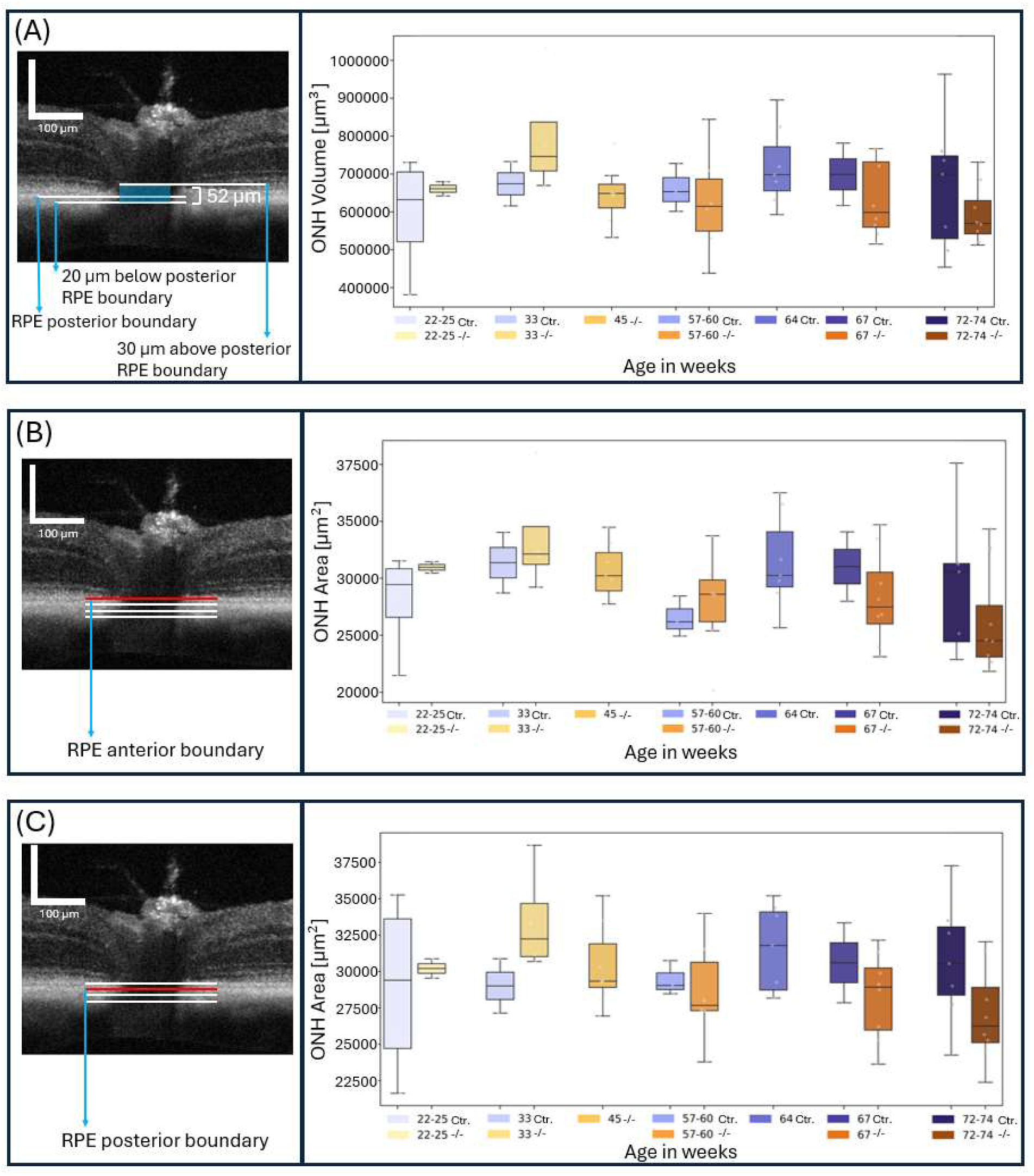
Comparison of ONH volume for -/- and control (Ctr.) mice of the SOD1 model. (A) Volume in a 52 micron range around the posterior RPE. (B) Area at the depth of the anterior RPE boundary. (C) Area at the depth of the posterior RPE boundary (C). Boxplots show the median (line), 25-75 percentile (boxes), interquartile range (whiskers), as well as all individual measurements.

## Discussion

The ONH is central to retinal structure and function. The data presented in this work, resulting from the analysis of ONH cross-sectional areas, 52-micron range volumes, and larger volumes in a 142-µm range (Supplementary Figure 3), clearly demonstrate that these ONH parameters depend on genotype, age, and sex. In the 5xFAD mouse model, similar characteristics were observed both for area measurements taken in different depths and for volume measurements. Notably, differences between tg and ntg 5xFAD models were observed, in particular at 9 months of age with a stronger visible difference for female tg animals. This finding aligns well with the more robust phenotype observed in female 5xFAD animals.^24^ Additionally, aging effects in 5xFAD animals were observed as a progressive decline in ONH volume over the lifespan of the animals.

Comparing multiple mouse models of AD yielded similar results for amyloid mouse models (APP/PS1 and 5xFAD) where larger ONH parameters were measured in tg mice than in ntg mice for APP/PS1 and 5xFAD animals. Given the genetic background of the mice, this difference between groups may be attributed to the amyloid pathology found in the models and aligns well with the findings of Cao and colleagues, who reported amyloid-beta accumulations in the ON of 5xFAD animals.^14^ In contrast, the PS19 mice, a model of tau pathology, did not display differences between tg and ntg animals. This hints towards a stronger influence of the amyloid pathology than the tau pathology on the ONH. Future studies with more animals of other models of amyloid and tau pathology will be needed to conclusively support this theory.

For the SOD1 mouse model dataset, ONH measurements presented a characteristic unlike that observed in the 5xFAD mice. A slight, yet non-significant age-dependence was observed but the effect was much weaker than in the 5xFAD mouse model. It should be noted that the most prominent volume changes for 5xFAD mice were observed between 3 and 9 months of age, with an initial increase of 6.7% from 3 to 5 months and then a decrease of 35% until 9 months of age. For older ages, the ONH parameters decreased rather slowly. This observation needs to be considered with caution as there are two factors potentially contributing to this quite different result. First, the SOD1 data has only been collected starting at the age of 5.5 months as compared to 3 months for the 5xFAD model. Second, the sample size and sampling density in the SOD1 study was low (n(ctr.) = 4, n(knockout) = 4), such that for a clear conclusion on the ONH changes in this model a larger cohort of animals will be necessary. Nevertheless, the applied method yields differing results for the SOD1 knockout and control mice and thus seems an interesting candidate to study changes of ONH in models of retinal degeneration. Another potential source of differences, is the degeneration of melanin reported for the model.^23^ The depth of the RPE proved to be the ideal depth for investigating the ONH area and volume in the studied mouse models because the signal-to-noise ratio was highest in this region and the segmentation was thus the most trustworthy (see Supplementary Figure 2). Notwithstanding the rather simple yet effective nature of the approach presented here, we believe that a quantitative analysis of the entire ONH structure is likely possible using more sophisticated segmentation methods. A more comprehensive analysis of the ONH geometry may yield a better understanding of the pathological changes in the ONH and ON and also prove useful for disease models in other rodents such as rats. Applying a similar segmentation could also be beneficial for the study of human ocular disease. In many studies such as for ONH drusen,^37^ myopia,^38^ and glaucoma research in non-human primates,^39^ manual segmentation is performed, leaving room for improvement via implementation of automatic algorithms such as the one presented here. However, it is important to note that ON anatomy for mice and humans/primates are not the same, as rodents do not possess a lamina cribrosa.

A target of future investigation could be the scleral canal of mouse models. Potential changes to this structure and the anterior portion of the ON could be assessed and yield more direct insight into pathologies affecting the ON. The scleral canal in glaucomatous humans has been imaged using spectral domain OCT, where canal expansion preceded RNFL defects,^40^ making the structure also interesting for mouse research. Albino animals, that lack the pigments obscuring this structure, would allow a direct in-vivo investigation of the SC.

Deteriorating retinal blood supply was reported for the ONH area in patients with AD, potentially caused by microvascular changes.^41^ Tsai and colleagues reported an increase of cup volume and cup to disc ratio as well as thinning of the disc rim in AD patients, indicating pathological changes to the ONH, related to the duration and severity of the disease in human patients.^42^ Additionally, larger optic cup to disk ratio combined with reduced ganglion cell axons, was also reported in a 2006 study by Meyer et al.^43^ Both of these findings align with the parameters measured for AD mouse models in this study and highlight a potential use of the ONH metrics as an ocular biomarker for AD in diagnostics. Given the unclear literature, with sometimes conflicting results, regarding standard retinal parameters such as retinal layer thickness and vascular parameters for the 5xFAD model^31, 44, 45, 46, 47^, the ONH potentially represents a more promising target. The analysis proposed here collectively assessed the vascular and neural components in the ONH. From the current data, we cannot untangle whether an observed change in area or volume was related to neural tissue, vasculature, or both. In our previous analysis of retinal vasculature in the 5xFAD mouse model, based on the same image data investigated here, no age-related changes were observed.^31^ Hence, it is more likely that the longitudinal ON area/volume decrease originated from a loss of neural tissue. Yet, since an influence of the vascular component cannot be excluded with certainty, further experiments will be required to untangle these two components of the ONH. Confirmation in models of vascular degeneration as well as correlative histology of the ONH and ON could reveal if the ONH analysis in this work can serve as a simple surrogate marker for axonal health or density measurements. Whether findings in the ONH precede RNFL loss would then be an exciting research question for targeted studies.

One limitation of the three analysis methods presented in this work is their dependence on the actual area scanned on the retina. For the mouse eye model assumed in our work,^19^ the scan range of the OCT system of 30° × 30° translated into a fixed field of view of 1mm × 1mm on the mouse retina. However, the field of view for a given scan angle depends on ocular parameters such as geometrical length, shape and refractive index of the various ocular structures (cornea, aqueous, crystalline lens, vitreous), which typically change throughout the life of a mouse.^48, 49^ They may also depend on genotype, sex, as well as pathological changes. Although axial eye length changes slow down in mice after the age of 60 days,^50, 51^ a very slow growth continues until at least 300 days of age.^51^ Of note: considerable variation in lens weight between strains has also been reported,^52^ as well as strong lens growth during development and slow growth until the end of life.^53^ Quantitative results between strains might thus not be uniform and change over time. Axial eye length increase has been measured for C57BL/6J mice and amounted to about 1.5% from 4-6 months of age and about 3% increase from 4 to 20 months of age. A substantial increase of 5% was measured from 2 to 4 months of age.^54^ The initial increase in volume/area measured from 3 to 5 months of age in the presented data for 5xFAD mice could therefore be an effect of aging related axial eye length increase. For later timepoints than 5 months, a change of 1.5% in axial length can be assumed. For a 30°×30° field of view, an increase in axial eye length of 1.5% translates to about 2.56% increase in area, a change much smaller than the observed changes for aging and pathology. Nevertheless, potential future refinements and improvements of our approach could include the measurement of eye length to correct for eye size changes during the mouse life.

The ONH is an anatomical feature easily accessible with most OCT systems, making an application of the presented segmentation broadly applicable. Segmentation was mostly based on intensity contrast alone (apart from the RPE segmentation), making the analysis a reasonable option for future studies and previously acquired datasets. Given the strong similarity in the observed behavior of volume to area measurements at the depth of the RPE, the investigation of areas alone might also be a reasonable option to assess ONH related changes. If depth dependent pathologies can be assumed, for example during longitudinal disease progression, segmenting areas in different imaging depths could potentially even provide additional insight into the investigated disease and its progression.

In summary, we presented a simple analysis approach for the quantification of ONH features in volumetric OCT data. Characteristic changes to the ONH were observed in several mouse models, suggesting the value of the ONH metrics as a potential biomarker for preclinical studies of ocular pathologies.

## Acknowledgements

The authors thank Sonja Reynoso-De-Leon, Christian Schönauer, Jasmin Rezek, and the team for their assistance with the mice. We also thank Eva Fuchs and Patrick Bilic for their support on animal welfare-related issues.

This project was supported by Scantox Neuro GmbH, FFG grant FO999900435, ERC Proof of Concept grant 101069344 OPTIMEYEZ, ERC Starting Grant 640396 OPTIMALZ, the Austrian Science Fund (FWF) grants I6092-B [doi:10.55776/I6092] and P35557 [doi:10.55776/P35557].

## Commercial Relationships Disclosure

G. Ladurner, None; M. Augustin, None; D. J. Harper, None; S. Worm, None; M. Varaka, None; L. May, None; Y. Patel, None; T. Rohrmoser, None; F. García-Ramírez, None; G. Garhöfer; None, M. Prokesch, Employment (Scantox Neuro GmbH); B. Baumann, None; and C. W. Merkle, None;

## References

1. Wang YX, Panda-Jonas S, Jonas JB. Optic nerve head anatomy in myopia and glaucoma, including parapapillary zones alpha, beta, gamma and delta: Histology and clinical features. Prog Retin Eye Res. Elsevier Ltd. 2021;83. doi:10.1016/j.preteyeres.2020.100933

2. Varadarajan SG, Huberman AD. Assembly and repair of eye-to-brain connections. Curr Opin Neurobiol. 2018;53:198–209. doi:10.1016/j.conb.2018.10.001

3. Cao S, Liao R, Fang C. Alterations in blood flow at the optic nerve head in patients with thyroid eye disease using optic coherence tomography angiography. Front Med (Lausanne). 2025;12. doi:10.3389/fmed.2025.1585907

4. Yoon J, Sung KR, Kim KE, Han HJ, Kim JM. Changes in optic nerve head microvasculature following disc hemorrhage absorption in glaucomatous eyes. Sci Rep. 2025;15(1). doi:10.1038/s41598-025-86460-7

5. Bata AM, Fondi K, Witkowska KJ, et al. Optic nerve head blood flow regulation during changes in arterial blood pressure in patients with primary open-angle glaucoma. Acta Ophthalmol. 2019;97(1):e36–e41. doi:10.1111/aos.13850

6. Rosario Hernandez M, O Pena JD, mary P. The Optic Nerve Head in Glaucomatous Optic Neuropathy. Arch Ophthalmol. 1997;115:389–395. http://archopht.jamanetwork.com/

7. Cao D, Yang D, Yu H, et al. Optic nerve head perfusion changes preceding peripapillary retinal nerve fibre layer thinning in preclinical diabetic retinopathy. Clin Exp Ophthalmol. 2019;47(2):219–225. doi:10.1111/ceo.13390

8. Şimşek M, Çitirik M, Özateş S, Şimşek T. Quantitative analysis of the optic nerve head parameters in patients with age-related macular degeneration. Turk J Med Sci. 2019;49(5):1518–1523. doi:10.3906/sag-1904-210

9. Wichrowska M, Wichrowski P, Kocięcki J. Morphological and Functional Assessment of the Optic Nerve Head and Retinal Ganglion Cells in Dry vs Chronically Treated Wet Age-Related Macular Degeneration. Clinical Ophthalmology. 2022;16:2373–2384. doi:10.2147/OPTH.S372626

10. Mirzaei N, Shi H, Oviatt M, et al. Alzheimer’s Retinopathy: Seeing Disease in the Eyes. Front Neurosci. Frontiers Media S.A. 2020;14. doi:10.3389/fnins.2020.00921

11. Kanaan NM, Pigino GF, Brady ST, Lazarov O, Binder LI, Morfini GA. Axonal degeneration in Alzheimer’s disease: When signaling abnormalities meet the axonal transport system. Exp Neurol. 2013;246:44–53. doi:10.1016/j.expneurol.2012.06.003

12. Sadun AA, Bassi CJ. Optic Nerve Damage in Alzheimer’s Disease. Ophthalmology. 1990;97(1):9–17. doi:10.1016/S0161-6420(90)32621-0

13. Tsai CS, Ritch R, Schwartz; Bernard, et al. Optic Nerve Head and Nerve Fiber Layer in Alzheimer’s Disease. doi:doi:10.1001/archopht.1991.01080020045040

14. Cao Q, Yang S, Wang X, et al. Transport of β-amyloid from brain to eye causes retinal degeneration in Alzheimer’s disease. Journal of Experimental Medicine. 2024;221(11). doi:10.1084/jem.20240386

15. Minakaran N, de Carvalho ER, Petzold A, Wong SH. Optical coherence tomography (OCT) in neuro-ophthalmology. Eye (Basingstoke). Springer Nature. 2021;35(1):17–32. doi:10.1038/s41433-020-01288-x

16. Shin HJ, Costello F. Imaging the optic nerve with optical coherence tomography. Eye (Basingstoke). Springer Nature. 2024;38(12):2365–2379. doi:10.1038/s41433-024-03165-3

17. Aumann S, Donner S, Fischer J, Müller F. Optical Coherence Tomography (OCT): Principle and Technical Realization. In: Bille JF, ed. High Resolution Imaging in Microscopy and Ophthalmology New Frontiers in Biomedical Optics. Springer; 2019:3. 10.1007/978-3-030-16638-0

18. Costello F, Rothenbuehler SP, Sibony PA, Hamann S, the Optic Disc Drusen Studies Consortium. Diagnosing Optic Disc Drusen in the Modern Imaging Era: A Practical Approach. Neuro-Ophthalmology. 2021;45(1):1–16. doi:10.1080/01658107.2020.1810286

19. Fialová S, Augustin M, Fischak C, et al. Posterior rat eye during acute intraocular pressure elevation studied using polarization sensitive optical coherence tomography. Biomed Opt Express. 2017;8(1):298. doi:10.1364/boe.8.000298

20. de Carlo TE, Romano A, Waheed NK, Duker JS. A review of optical coherence tomography angiography (OCTA). Int J Retina Vitreous. BioMed Central Ltd. 2015;1(1). doi:10.1186/s40942-015-0005-8

21. Steiner S, Schwarzhans F, Desissaire S, et al. Birefringent Properties of the Peripapillary Retinal Nerve Fiber Layer in Healthy and Glaucoma Subjects Analyzed by Polarization-Sensitive OCT. Invest Ophthalmol Vis Sci. 2022;63(12). doi:10.1167/iovs.63.12.8

22. Desissaire S, Pollreisz A, Sedova A, et al. Analysis of retinal nerve fiber layer birefringence in patients with glaucoma and diabetic retinopathy by polarization sensitive OCT. Biomed Opt Express. 2020;11(10):5488. doi:10.1364/boe.402475

23. Merkle CW, Augustin M, Harper DJ, Glösmann M, Baumann B. Degeneration of Melanin-Containing Structures Observed Longitudinally in the Eyes of SOD1−/− Mice Using Intensity, Polarization, and Spectroscopic OCT. Transl Vis Sci Technol. 2022;11(10). doi:10.1167/tvst.11.10.28

24. Forner S, Kawauchi S, Balderrama-Gutierrez G, et al. Systematic phenotyping and characterization of the 5xFAD mouse model of Alzheimer’s disease. Sci Data. 2021;8(1). doi:10.1038/s41597-021-01054-y

25. Imamura Y, Noda S, Hashizume K, et al. Drusen, choroidal neovascularization, and retinal pigment epithelium dysfunction in SOD1-deficient mice: A model of age-related macular degeneration. PNAS. 2006;103 (30):11282–11287. 10.1073/pnas.0602131103

26. Forner S, Kawauchi S, Balderrama-Gutierrez G, et al. Systematic phenotyping and characterization of the 5xFAD mouse model of Alzheimer’s disease. Sci Data. 2021;8(1). doi:10.1038/s41597-021-01054-y

27. Zhong MZ, Peng T, Duarte ML, Wang M, Cai D. Updates on mouse models of Alzheimer’s disease. Mol Neurodegener. 2024;19(1). doi:10.1186/s13024-024-00712-0

28. Hashizume K, Hirasawa M, Imamura Y, et al. Retinal dysfunction and progressive retinal cell death in SOD1-deficient mice. American Journal of Pathology. 2008;172(5):1325–1331. doi:10.2353/ajpath.2008.070730

29. Ladurner G, Merkle CW, May L, et al. Longitudinal investigation of spatial memory and retinal parameters in a 5xFAD model of Alzheimer’s disease reveals differences dependent on genotype and sex. Biomed Opt Express. 2025;17(1):405. doi:10.1364/boe.579020

30. Fialová S, Augustin M, Glösmann M, et al. Polarization properties of single layers in the posterior eyes of mice and rats investigated using high resolution polarization sensitive optical coherence tomography. Biomed Opt Express. 2016;7(4):1479. doi:10.1364/boe.7.001479

31. Ladurner G, Harper DJ, May L, et al. Comparative Investigation of the Retinal Phenotype of Three Mouse Models of Alzheimer’s Disease With Optical Coherence Tomography. Invest Ophthalmol Vis Sci. 2025;66(14). doi:10.1167/iovs.66.14.35

32. Augustin M, Fialová S, Himmel T, et al. Multi-functional OCT enables longitudinal study of retinal changes in a VLDLR knockout mouse model. PLoS One. 2016;11(10). doi:10.1371/journal.pone.0164419

33. Götzinger E, Pircher M, Geitzenauer W, et al. Retinal pigment epithelium segmentation by polarization sensitive optical coherence tomography. Opt Express. 2008;16, Nr. 21(170):16410–16422. doi:10.1364/oe.16.016410

34. Baumann B, Baumann SO, Konegger T, et al. Polarization sensitive optical coherence tomography of melanin provides intrinsic contrast based on depolarization. Biomed Opt Express. 2012;254(3):1670–1683.

35. Augustin M, Harper DJ, Merkle CW, Hitzenberger CK, Baumann B. Segmentation of Retinal Layers in OCT Images of the Mouse Eye Utilizing Polarization Contrast. In: Lecture Notes in Computer Science (Including Subseries Lecture Notes in Artificial Intelligence and Lecture Notes in Bioinformatics). 11039 LNCS. Springer Verlag; 2018:310–318. doi:10.1007/978-3-030-00949-6_37

36. Augustin M, Harper DJ, Merkle CW, Glösmann M, Hitzenberger CK, Baumann B. Optical coherence tomography findings in the retinas of SOD1 knockout mice. Transl Vis Sci Technol. 2020;9(4):1–12. doi:10.1167/tvst.9.4.15

37. Tsikata E, Vercellin Verticchio AC, Falkenstein I, et al. Volumetric Measurement of Optic Nerve Head Drusen Using Swept-Source Optical Coherence Tomography. J Glaucoma. 2017;26(9):798–804. doi:10.1097/IJG.0000000000000707

38. Jeoung JW, Yang H, Gardiner S, et al. Optical Coherence Tomography Optic Nerve Head Morphology in Myopia I: Implications of Anterior Scleral Canal Opening Versus Bruch Membrane Opening Offset. Am J Ophthalmol. 2020;218:105–119. doi:10.1016/j.ajo.2020.05.015

39. Yang H, Qi J, Hardin C, et al. Spectral-domain optical coherence tomography enhanced depth imaging of the normal and glaucomatous nonhuman primate optic nerve head. Invest Ophthalmol Vis Sci. 2012;53(1):394–405. doi:10.1167/iovs.11-8244

40. Sawada Y, Ishikawa M, Iwase T, Yoshitomi T, Araie M. Clinical assessment of scleral canal expansion in glaucoma using spectral domain optical coherence tomography. Invest Ophthalmol Vis Sci. 2020;61(7):1965.

41. Bambo MP, Garcia-Martin E, Gutierrez-Ruiz F, et al. Analysis of optic disk color changes in Alzheimer’s disease: A potential new biomarker. Clin Neurol Neurosurg. 2015;132:68–73. doi:10.1016/j.clineuro.2015.02.016

42. Tsai CS, Ritch R, Schwartz B, et al. Optic Nerve Head and Nerve Fiber Layer in Alzheimer’s Disease. JAMA Ophthalmol. 1991;109, No.2:199–204. doi:doi:10.1001/archopht.1991.01080020045040

43. Danesh-Meyer H V, Birch ; H, Ku YF, Carroll ; S, Gamble G. Reduction of optic nerve fibers in patients with Alzheimer disease identified by laser imaging From the Department of Ophthalmology (2006;(67):1852–1854. 10.1212/01.wnl.0000244490.07925.8

44. Lim JKH, Li QX, He Z, et al. Retinal Functional and Structural Changes in the 5xFAD Mouse Model of Alzheimer’s Disease. Front Neurosci. 2020;14. doi:10.3389/fnins.2020.00862

45. Kim TH, Son T, Klatt D, Yao X. Concurrent OCT and OCT angiography of retinal neurovascular degeneration in the 5XFAD Alzheimer’s disease mice. Neurophotonics. 2021;8(03). doi:10.1117/1.nph.8.3.035002

46. Matei N, Leahy S, Blair NP, Burford J, Rahimi M, Shahidi M. Retinal Vascular Physiology Biomarkers in a 5XFAD Mouse Model of Alzheimer’s Disease. Cells. 2022;11(15). doi:10.3390/cells11152413

47. Lynn SA, Pandi SPS, Sanchez-Bretano A, et al. A longitudinal study of the 5xFAD mouse retina delineates Amyloid beta (Aβ)-mediated retinal pathology from age-related changes. Alzheimer’s Research and Therapy . 2025;17(1). doi:10.1186/s13195-025-01784-w

48. Jiang M, Wu PC, Fini ME, et al. Single-shot dimension measurements of the mouse eye using SD-OCT. Ophthalmic Surgery Lasers and Imaging. 2012;43(3):252–256. doi:10.3928/15428877-20120308-04

49. Wisard J, Chrenek MA, Wright C, et al. Non-contact measurement of linear external dimensions of the mouse eye. J Neurosci Methods. 2010;187(2):156–166. doi:10.1016/j.jneumeth.2010.01.006

50. Zhou X, Shen M, Xie J, et al. The development of the refractive status and ocular growth in C57BL/6 mice. Invest Ophthalmol Vis Sci. 2008;49(12):5208–5214. doi:10.1167/iovs.07-1545

51. Tkatchenko T V., Shen Y, Tkatchenko A V. Analysis of postnatal eye development in the mouse with high-resolution small animal magnetic resonance imaging. Invest Ophthalmol Vis Sci. 2010;51(1):21–27. doi:10.1167/iovs.08-2767

52. Zhou G, Williams WR. Mouse Models for the Analysis of Myopia: An Analysis of Variation in Eye Size of Adult Mice. Optometry and Vision Science. 1999;76(6):408–418.

53. Painter T, Ou C, Gong X, Xia CH. Longitudinal study of microphthalmia in connexin 50 knockout mice using spectral-domain optical coherence tomography. Frontiers in Ophthalmology. 2024;4. doi:10.3389/fopht.2024.1387961

54. Chou TH, Kocaoglu OP, Borja D, et al. Postnatal elongation of eye size in DBA/2J mice compared with C57BL/6J mice: In vivo analysis with whole-eye OCT. Invest Ophthalmol Vis Sci. 2011;52(6):3604–3612. doi:10.1167/iovs.10-6340

