## Supplementary Material for "Exploration of Structural Optic Nerve Changes in Mouse Models of Retinal and Neuronal Degeneration with Optical Coherence Tomography"

**Supplementary Table 1**: Animal number for each imaging timepoint split by genotype and sex for the 5xFAD mouse model.

| Age in Weeks | | 12 | | 20 | | 24 | | 36 | | 96-104 | |
|---|---|---|---|---|---|---|---|---|---|---|---|
| Tg 5xFAD | | 32 | | 24 | | 24 | | 14 | | / | |
| Ntg 5xFAD | | 32 | | 24 | | 24 | | 13 | | 6 | |
| Tg Female | Tg Male | 16 | 16 | 12 | 12 | 12 | 12 | 8 | 6 | / | / |
| Ntg Female | Ntg Male | 16 | 16 | 12 | 12 | 12 | 12 | 8 | 5 | 3 | 3 |

**Supplementary Table 2**: Animal number for each model split by genotype for the 5xFAD, PS19 and APP/PS1 mouse model.

| | 5xFAD | PS19 | APP/PS1 |
|---|---|---|---|
| Age in Weeks | 36 | 39 | 51-54 |
| Tg | 14 | 3 | 4 |
| Ntg | 13 | 3 | 3 |

Supplementary Table 3: Animal number for each imaging timepoint split by genotype and sex for the 5xFAD mouse model.

| Age in Weeks | 22-25 | 33 | 45 | 57-60 | 64 | 67 | 72-74 |
|---|---|---|---|---|---|---|---|
| SOD1 +/+ | 2 | 1 | / | 2 | 4 | 1 | 4 |
| SOD1 Control | 1 | 2 | 4 | 3 | / | 4 | 4 |

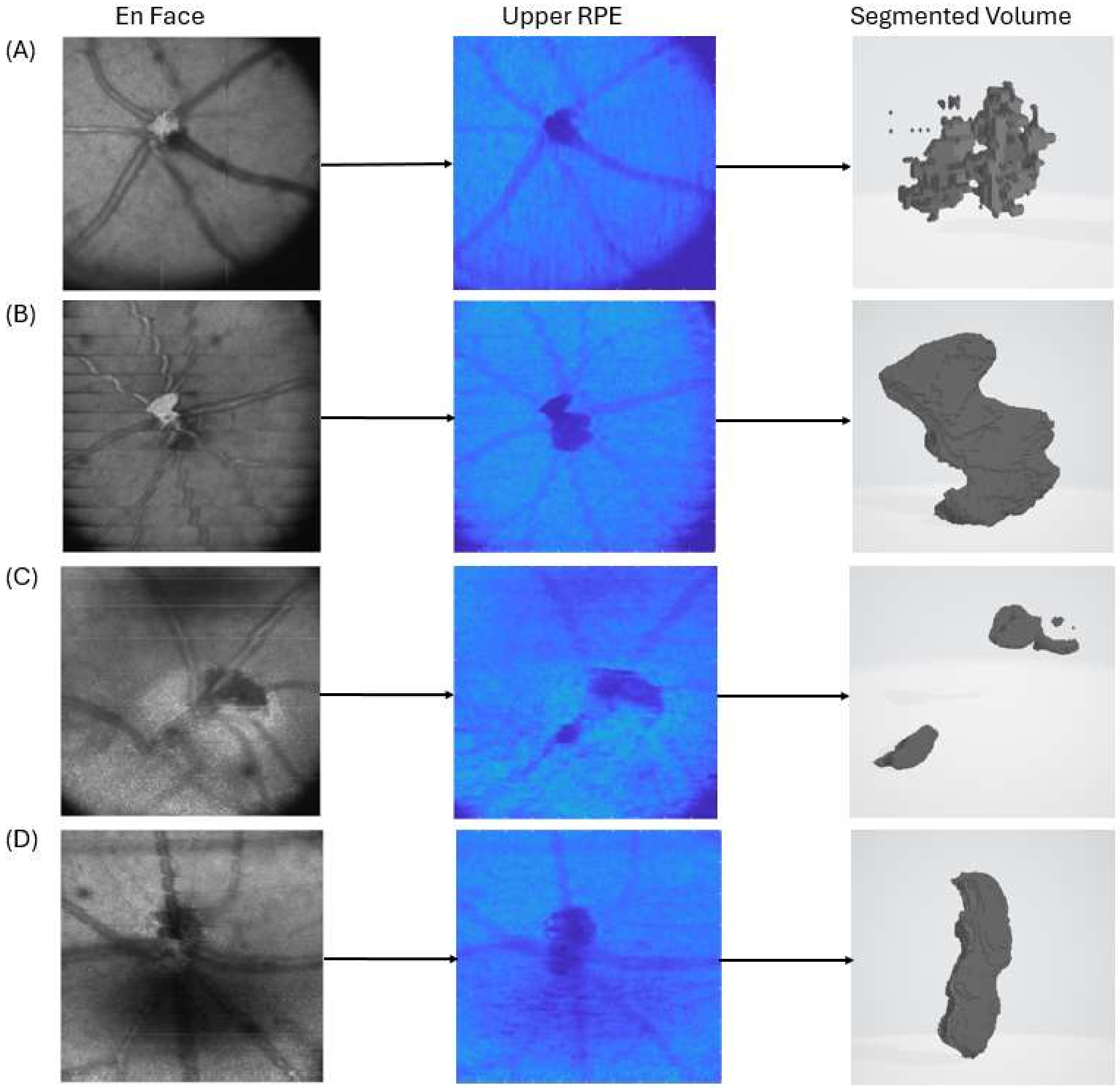


**Supplementary Figure 1**: Scans excluded from the analysis and the resulting intensity in the anterior RPE, as well as the resulting volume. Strong vignetting (A), strong breathing motion (B), obscured ONH (C), ONH segmentation hinder due to bad signal (D).

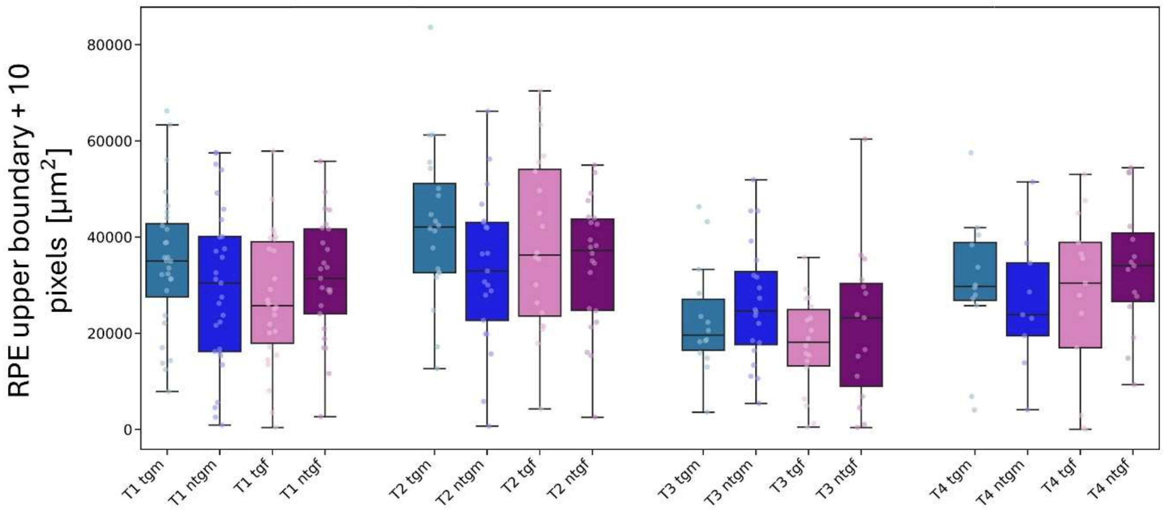


**Supplementary Figure 2**: Resulting area values for the segmentation placed 20 μm above the RPE with $TI_{1.7}$.

*Quantification of Extended Volumes of the ONH space*

To assess the three-dimensional structure of the ONH, we quantitatively analyzed extended volumes composed of segmented areas stacked on each other. The posterior RPE boundary was again chosen as a reference and all areas within a 142-μm deep range extending from 35 pixels above to 35 pixels below the posterior RPE were segmented using the method described for areas with $TI_{2.5}$ as a threshold. Then, the average slice area was calculated by averaging over all 71 segmented areas. By multiplying this number with the height of 142μm, volumes are calculated. Figure 1E shows the resulting volumes that encompass 71 pixels in the z-direction. This specific range was chosen as above and below this 71-pixel range, the segmentation sometimes failed due to low intensity contrast.

*Results for Extended Volumes of the ONH space*

To gain an understanding of structural changes in a larger portion of the ONH, we evaluated all large ONH volumes in the 71-pixel-deep range. An example of the analyzed region and the resulting volume are shown in a B-scan in Supplementary Figure 3A. Volumes for 5xFAD mice are shown in Supplementary Figure 3(B-C).

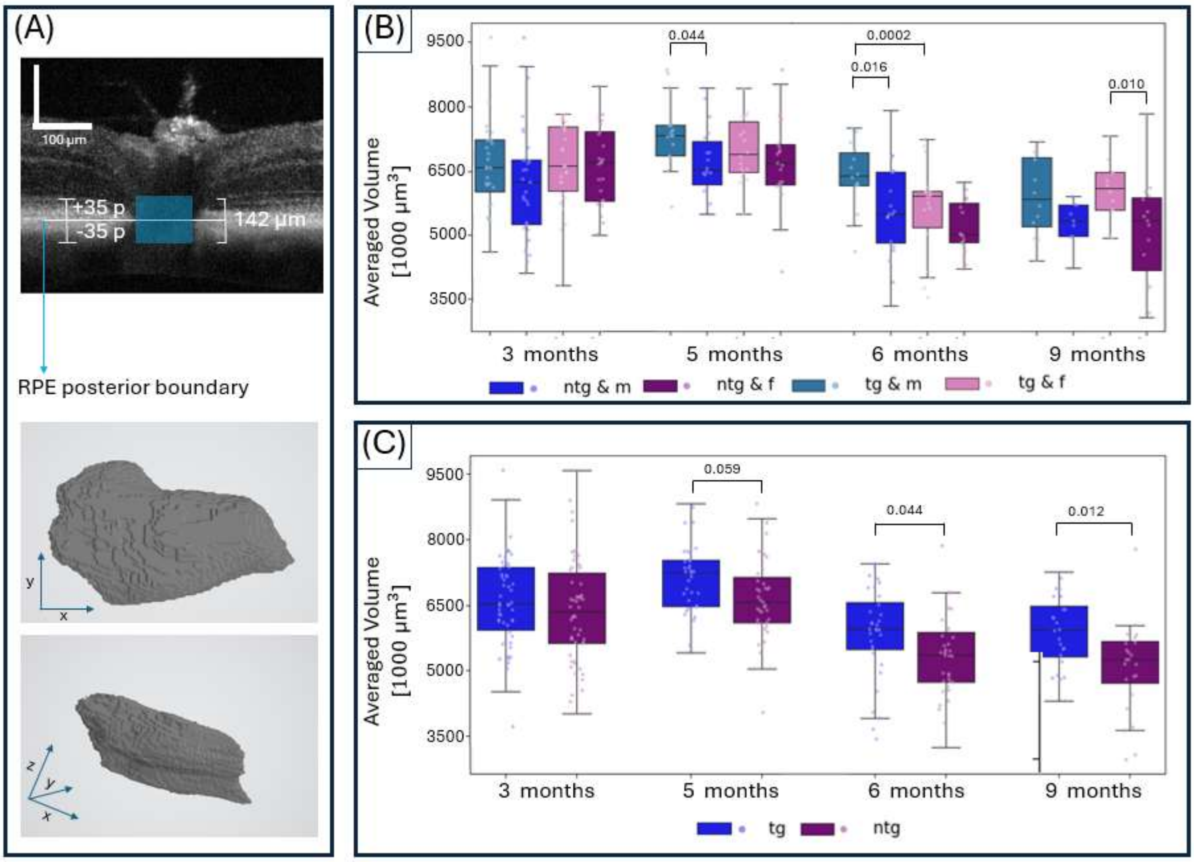


**Supplementary Figure 3**. Longitudinal development of averaged ONH volumes for 5xFAD mice evaluated for a 71-pixel range around the posterior RPE boundary. (A) Visualization of the evaluated depth range in a B-scan and exemplary volume renderings of the ONH structure. (B) Longitudinal ONH volume stratified for sex and genotype. (C) Longitudinal ONH volume for ntg and tg animals after pooling female and male subjects. Boxplots show the median (line), 25-75 percentile (boxes), interquartile range (whiskers), as well as all individual measurements.

Supplementary Figure 3B shows the development of the volume for tgm tgf, ntgm and ntgf animals over the 6-month study course. Volumes increased by about 11.5% from 3 to 5 months of age followed by a decrease by 21% from 5 to 9 months of age. Similar longitudinal characteristics to the area measurements were observed. Significant volume differences between groups were found between tgm and ntgm mice at 5 and 6 months of age, with (54,156µm$^3$±5,970µm$^3$) for tgm and (49,518µm$^3$±6,118µm$^3$) for ntgm at 5 months ($p=0.044$). At 6 months of age, tgm (47,730µm$^3$±6,187µm$^3$) and ntgm µ(41,849µm$^3$±8,186µm$^3$) groups were significantly different ($p=0.016$). Values for tgm and ntgm animals differ by 8.5% at 5 months and by 13% at 6 months. For tgf mice at 9 months of age, an averaged volume of 44,295µm$^3$±5,144µm$^3$ was measured, significantly larger than the volume for ntgf mice (37,495µm$^3$±9,300µm$^3$). All measured mean values and standard deviations are listed in Supplementary Tables 4 and 5.

Tg animals display larger volumes than ntg mice. For tg animals averaged volumes of 42,232µm$^3$±4,645µm$^3$, 3,5618µm$^3$±6,261µm$^3$ and 35,042µm$^3$±4,748µm$^3$ were measured for 5, 6 and 9 months of age, respectively (Supplementary Figure 3C). For ntg animals the analysis resulted in lower volumes of 39,669µm$^3$±5,580µm$^3$, 32,503µm$^3$±5,583µm$^3$ and 30,397µm$^3$±6,093µm$^3$ for 5, 6 and 9 months of age respectively. These volume data correspond to a difference between tg and ntg animals of 6.1% at 5 months ($p=0.059$), 8.8% at 6 months ($p=0.044$) and 13.4% at 9 months ($p=0.012$).

**Supplementary Table 4**: Mean and standard deviation for measured areas and volumes for male and female tg and ntg 5xFAD animals.

| 3 Months of Age | | | | | | | |
|---|---|---|---|---|---|---|---|
| Area Location | | RPE anterior boundary | RPE posterior boundary | 20 µm below posterior RPE | 40 µm below posterior RPE | Volume | Volume (26 pixels) |
| Mean Values | tgf | 27010.38 | 26829.71 | 30256.83 | 30201.54 | 48878.33 | 831721.88 |
| | ntgf | 26234.30 | 26323.30 | 30328.50 | 30010.60 | 48580.99 | 862251.75 |
| | tgm | 26919.82 | 26977.32 | 30719.64 | 31664.46 | 49152.04 | 857122.77 |
| | ntgm | 25089.66 | 25339.14 | 29173.97 | 27744.83 | 46254.16 | 753941.81 |
| Standard Deviation | tgf | 4420.06 | 3603.82 | 4133.87 | 5340.17 | 7436.47 | 201611.09 |
| | ntgf | 4601.21 | 3266.03 | 3924.91 | 6883.36 | 7152.80 | 222952.21 |
| | tgm | 4803.48 | 4049.69 | 4024.58 | 4973.50 | 8144.47 | 206897.75 |
| | ntgm | 5890.89 | 4867.23 | 5832.53 | 7546.00 | 10001.25 | 234161.45 |

| 5 Months of Age | | | | | | | |
|---|---|---|---|---|---|---|---|
| Area Location | | RPE anterior boundary | RPE posterior boundary | 20 µm below posterior RPE | 40 µm below posterior RPE | Volume | Volume (26 pixels) |
| Mean Values | tgf | 27545.25 | 26866.50 | 31082.25 | 33153.5 | 51492.49 | 987422.92 |
| | ntgf | 27498.18 | 26529.43 | 30982.39 | 31794.88636 | 49644.37 | 869001.79 |
| | tgm | 28864.50 | 27814.75 | 32595.00 | 33614.5 | 54165.16 | 985152.88 |
| | ntgm | 27660.95 | 27756.19 | 31467.86 | 29019.52381 | 49518.74 | 846384.03 |
| Standard Deviation | tgf | 3111.39 | 2578.06 | 3651.61 | 6459.77 | 5496.27 | 211147.68 |
| | ntgf | 4129.87 | 4277.49 | 4359.82 | 4651.61 | 7785.80 | 220382.95 |
| | tgm | 5013.08 | 4457.05 | 4682.37 | 5002.34 | 5970.20 | 160534.11 |
| | ntgm | 5172.75 | 4198.71 | 5692.74 | 7115.22 | 6118.17 | 144561.67 |

| 6 Months of Age | | | | | | | |
|---|---|---|---|---|---|---|---|
| Area Location | | RPE anterior boundary | RPE posterior boundary | 20 µm below posterior RPE | 40 µm below posterior RPE | Volume | Volume (26 pixels) |

| | | | | | | | |
|---|---|---|---|---|---|---|---|
| Mean Values | tgf | 20668.00 | 20950.46 | 21925.49 | 18535.97 | 42028.60 | 701237.26 |
| | ntgf | 19649.89 | 19920.30 | 21073.87 | 17725.91 | 39003.22 | 678769.37 |
| | tgm | 23628.10 | 23458.36 | 25620.09 | 24825.54 | 47730.56 | 814231.69 |
| | ntgm | 20688.29 | 20454.30 | 22998.69 | 20007.66 | 41849.99 | 745048.31 |

| | | | | | | | |
|---|---|---|---|---|---|---|---|
| Standard Deviation | tgf | 4715.03 | 3422.14 | 5061.82 | 6445.02 | 8208.93 | 262706.72 |
| | ntgf | 2703.96 | 2216.19 | 3191.71 | 5506.91 | 4733.36 | 193321.24 |
| | tgm | 3470.37 | 2838.22 | 3124.04 | 3762.96 | 6187.09 | 228472.66 |
| | ntgm | 4715.03 | 3422.14 | 5061.82 | 6445.02 | 8186.20 | 221498.00 |

9 Months of Age

| Area Location | | RPE anterior boundary | RPE posterior boundary | 20 µm below posterior RPE | 40 µm below posterior RPE | Volume | Volume (26 pixels) |
|---|---|---|---|---|---|---|---|
| Mean Values | tgf | 20204.92 | 24493.62 | 25054.55 | 24308.65 | 44295.69 | 812456.22 |
| | ntgf | 19692.96 | 19445.40 | 22403.83 | 21442.78 | 37495.33 | 552859.56 |
| | tgm | 22340.91 | 21955.61 | 23394.60 | 21774.12 | 43310.88 | 718471.63 |
| | ntgm | 20300.96 | 19672.36 | 23394.60 | 21774.12 | 38833.24 | 579073.08 |

| | | | | | | | |
|---|---|---|---|---|---|---|---|
| Standard Deviation | tgf | 6763.09 | 11244.62 | 4487.45 | 6069.68 | 5145.57 | 191894.39 |
| | ntgf | 5143.30 | 4723.08 | 4659.84 | 3761.48 | 9300.41 | 185799.35 |
| | tgm | 2878.61 | 2862.34 | 3501.32 | 5895.31 | 6831.51 | 173693.64 |
| | ntgm | 2188.83 | 1316.08 | 2332.70 | 2476.67 | 3767.07 | 47366.01 |

**Supplementary Table 5**: Mean and standard deviation for measured areas and volumes for tg and ntg 5xFAD animals.

3 Months of Age

| Area Location | | RPE anterior boundary | RPE posterior boundary | 20 µm below posterior RPE | 40 µm below posterior RPE | Volume | Volume (26 pixels) |
|---|---|---|---|---|---|---|---|
| Mean Values | tg | 26963.43 | 26906.25 | 30496.81 | 30960.09 | 39216.20 | 844892.71 |
| | ntg | 25619.58 | 25794.77 | 29708.47 | 28793.80 | 37865.12 | 804085.30 |

| | | | | | | | |
|---|---|---|---|---|---|---|---|
| Standard Deviation | tg | 4579.52 | 3806.10 | 4045.58 | 5157.25 | 6191.42 | 202840.88 |
| | ntg | 5315.29 | 4194.17 | 5028.47 | 7269.01 | 7037.40 | 233341.96 |

| 5 Months of Age |
| --- |

| Area Location | | RPE anterior boundary | RPE posterior boundary | 20 µm below posterior RPE | 40 µm below posterior RPE | Volume | Volume (26 pixels) |
| --- | --- | --- | --- | --- | --- | --- | --- |
| Mean Values | tg | 28204.88 | 27340.63 | 31838.63 | 33384 | 42232.52 | 986470.97 |
| | ntg | 27577.67 | 27128.55 | 31219.48 | 30439.47674 | 39669.11 | 858562.82 |
| Standard Deviation | tg | 4172.03 | 3625.81 | 4214.76 | 5707.42 | 4645.15 | 186974.12 |
| | ntg | 4612.62 | 4234.18 | 4999.63 | 6074.29 | 5580.23 | 187195.25 |

| 6 Months of Age |
| --- |

| Area Location | | RPE anterior boundary | RPE posterior boundary | 20 µm below posterior RPE | 40 µm below posterior RPE | Volume | Volume (26 pixels) |
| --- | --- | --- | --- | --- | --- | --- | --- |
| Mean Values | tg | 21963.04 | 22047.67 | 23541.88 | 21287.66 | 35618.57 | 751457.01 |
| | ntg | 20243.26 | 20225.44 | 22173.76 | 19029.77 | 32503.96 | 717432.08 |
| Standard Deviation | tg | 4412.24 | 3376.10 | 4648.77 | 6226.26 | 6261.81 | 251155.56 |
| | ntg | 3419.54 | 3062.43 | 4312.34 | 5790.05 | 5583.83 | 209958.40 |

| 9 Months of Age |
| --- |

| Area Location | | RPE anterior boundary | RPE posterior boundary | 20 µm below posterior RPE | 40 µm below posterior RPE | Volume | Volume (26 pixels) |
| --- | --- | --- | --- | --- | --- | --- | --- |
| Mean Values | tg | 21230.20 | 23275.37 | 24707.29 | 24648.41 | 35042.63 | 767343.62 |
| | ntg | 19920.96 | 19530.51 | 22775.37 | 21567.03 | 30397.64 | 562689.63 |
| Standard Deviation | tg | 5277.67 | 8285.57 | 3977.88 | 5871.97 | 4748.80 | 185838.68 |
| | ntg | 4225.99 | 3767.43 | 3917.91 | 3282.17 | 6093.90 | 148193.92 |

**Supplementary Table 6**: Mean and standard deviation for measured volumes for ntg 5xFAD animals.

| | | 3 Months of Age | 5 Months of Age | 6 Months of Age | 9 Months of Age | 22-24 Months of Age |
| --- | --- | --- | --- | --- | --- | --- |
| Area Location | | Volume (26 pixels) | Volume (26 pixels) | Volume (26 pixels) | Volume (26 pixels) | Volume (26 pixels) |
| Mean Values | female | 862251.75 | 869001.79 | 678769.37 | 552859.56 | 423478.80 |

| | | | | | | |
|---|---|---|---|---|---|---|
| | male | 753941.81 | 846384.03 | 745048.31 | 579073.08 | 564541.41 |
| | all | 804085.30 | 858562.82 | 717432.08 | 562689.63 | 508116.37 |

| | | | | | | |
|---|---|---|---|---|---|---|
| Standard Deviation | female | 222952.21 | 220382.95 | 193321.24 | 185799.35 | 114344.87 |
| | male | 234161.45 | 144561.6 | 221498.00 | 173693.64 | 233606.38 |
| | all | 233341.96 | 187195.25 | 209958.40 | 148193.92 | 199955.76 |

**Supplementary Table 7**: Mean and standard deviation for measured areas and volumes for ntg and tg animals of 5xFAD, PS19 and APP/PS1 mouse lines.

| | 26 pixel volume [µm] | | Posterior RPE Area [µm] | | Anterior RPE Area [µm] | |
|---|---|---|---|---|---|---|
| | Average | Standard Deviation | Average | Standard Deviation | Average | Standard Deviation |
| 5xFAD tg | 844892.7 | 202840.9 | 26906.3 | 3806.1 | 26963.4 | 4579.5 |
| 5xFAD ntg | 804085.3 | 233342.0 | 25794.8 | 4194.2 | 25619.6 | 5315.3 |
| PS19 tg | 516001.7 | 107125.3 | 21011.3 | 2876.8 | 19775.3 | 4137.1 |
| PS19 ntg | 573655.0 | 103934.2 | 22161.3 | 2213.3 | 20791.3 | 4607.4 |
| APP/PS1 tg | 848672.5 | 68907.9 | 38352.5 | 3094.3 | 37041.9 | 2857.5 |
| APP/PS1 ntg | 738683.3 | 157625.6 | 33038.3 | 5368.9 | 32397.5 | 4995.5 |

**Supplementary Table 8**: P-values for the comparisons between different AD models.

P<0.05 = green, yellow 0.05<p<0.15, red p>0.15.

| | 26 Pixel Volume | | | | | |
|---|---|---|---|---|---|---|
| | 5xFAD tg | 5xFAD ntg | PS19 tg | PS19 ntg | APP/PS1 tg | APP/PS1 ntg |
| 5xFAD tg | | 9.39E-06 | 0.05850 | 0.39950 | 1.55E-06 | 0.01750 |
| 5xFAD ntg | 9.39E-06 | | 0.19765 | 0.01740 | 5.10E-13 | 3.23E-07 |
| PS19 tg | 0.05850 | 0.19765 | | 0.36643 | 4.53E-07 | 0.00086 |
| PS19 ntg | 0.39950 | 0.01740 | 0.36643 | | 0.00002 | 0.01463 |
| APP/PS1 tg | 1.55E-06 | 5.10E-13 | 4.53E-07 | 0.00002 | | 0.06888 |
| APP/PS1 ntg | 0.01750 | 3.23E-07 | 0.00086 | 0.01463 | 0.06888 | |

| | Posterior RPE Area | | | | | |
|---|---|---|---|---|---|---|
| | 5xFAD tg | 5xFAD ntg | PS19 tg | PS19 ntg | APP/PS1 tg | APP/PS1 ntg |
| 5xFAD tg | | 0.001125 | 0.239 | 0.692 | 0 | 1.09022E-11 |
| 5xFAD ntg | 0.001125 | | 0.34085 | 0.071591 | 0 | 1.11022E-15 |
| PS19 tg | 0.239 | 0.3408462 | | 0.503786 | 0 | 2.06544E-10 |
| PS19 ntg | 0.692 | 0.0715909 | 0.50379 | | 6.6612E-16 | 3.83688E-09 |
| APP/PS1 tg | 0 | 0 | 0 | 6.66E-16 | | 0.0009 |
| APP/PS1 ntg | 1.09E-11 | 1.11E-15 | 2.1E-10 | 3.84E-09 | 0.0009 | |

| | Anterior RPE Area | | | | | |
|---|---|---|---|---|---|---|
| | 5xFAD tg | 5xFAD ntg | PS19 tg | PS19 ntg | APP/PS1 tg | APP/PS1 ntg |
| 5xFAD tg | | 0.0045 | 0.0255 | 0.118375 | 0 | 2.43518E-08 |
| 5xFAD ntg | 0.0045 | | 0.8293 | 0.667607 | 0 | 3.90078E-12 |
| PS19 tg | 0.0255 | 0.8293 | | 0.664962 | 7.494E-15 | 3.16849E-09 |
| PS19 ntg | 0.118375 | 0.6676071 | 0.66496 | | 7.3275E-14 | 2.52093E-08 |
| APP/PS1 tg | 0 | 0 | 7.5E-15 | 7.33E-14 | | 0.01155 |
| APP/PS1 ntg | 2.44E-08 | 3.901E-12 | 3.2E-09 | 2.52E-08 | 0.01155 | |

**Supplementary Table 9**: Mean and standard deviation for measured areas and volumes for -/- and control SOD1 animals.

| | | 22-25 Weeks | 33 Weeks | 45 Weeks | 57-60 Weeks | 64 Weeks | 67 Weeks | 72-74 Weeks |
|---|---|---|---|---|---|---|---|---|
| Area Location | | Volume (26 pixels) | Volume (26 pixels) | Volume (26 pixels) | Volume (26 pixels) | Volume (26 pixels) | Volume (26 pixels) | Volume (26 pixels) |
| Mean Values | Knockout (-/-) | 594292.50 | 674445.00 | | 660826.67 | 720392.86 | 699350.00 | 667418.57 |
| | Control | 661000.00 | 798805.00 | 647067.14 | 625135.00 | | 634197.50 | 593578.75 |
| Standard Deviation | Knockout (-/-) | 158312.57 | 82766.85 | | 63574.42 | 106423.30 | 116347.35 | 176767.10 |
| | Control | 26318.51 | 161054.24 | 79866.43 | 140938.97 | | 102067.42 | 78154.63 |

| 22-25 Weeks | 33 Weeks | 45 Weeks | 57-60 Weeks | 64 Weeks | 67 Weeks | 72-74 Weeks |
|---|---|---|---|---|---|---|

| Area Location | | RPE posterior boundary | RPE posterior boundary | RPE posterior boundary | RPE posterior boundary | RPE posterior boundary | RPE posterior boundary | RPE posterior boundary |
|---|---|---|---|---|---|---|---|---|
| Mean Values | Knockout (-/-) | 28928.75 | 29015.00 | | 29426.67 | 31546.43 | 30602.50 | 30706.43 |
| | Control | 30210.00 | 33460.00 | 30440.71 | 28660.83 | | 28293.75 | 27045.00 |
| Standard Deviation | Knockout (-/-) | 6340.14 | 2644.58 | | 1185.75 | 3009.20 | 3878.48 | 4238.19 |
| | Control | 940.45 | 3665.87 | 2901.74 | 3584.84 | | 2991.03 | 3337.28 |
| | | 22-25 Weeks | 33 Weeks | 45 Weeks | 57-60 Weeks | 64 Weeks | 67 Weeks | 72-74 Weeks |

| Area Location | | RPE anterior boundary | RPE anterior boundary | RPE anterior boundary | RPE anterior boundary | RPE anterior boundary | RPE anterior boundary | RPE anterior boundary |
|---|---|---|---|---|---|---|---|---|
| Mean Values | Knockout (-/-) | 27978.75 | 31380.00 | | 26521.67 | 31446.43 | 31042.50 | 29290.71 |
| | Control | 30962.50 | 33636.25 | 30687.86 | 27792.50 | | 28316.25 | 26210.63 |
| Standard Deviation | Knockout (-/-) | 4546.30 | 3768.88 | | 1780.82 | 4244.44 | 4309.82 | 6008.00 |
| | Control | 703.57 | 5132.32 | 2514.19 | 4625.88 | | 4139.41 | 4686.77 |

**Supplementary Table 10**: Percentage of excluded scans for each measurement timepoint for 5xFAD, PS19 and APP/PS1 animals.

| Timepoint [weeks/months] | 12/3 | 20/5 | 24/6 | 36/9 | 39/9.5 | 51-54/11-12.5 |
|---|---|---|---|---|---|---|
| 5xFAD | 11.2% | 12.3% | 16.2% | 10.7% | / | / |
| APP/PS1 | / | / | / | / | / | 13.3% |
| PS19 | / | / | / | / | 0% | / |

**Supplementary Table 11**: Percentage of excluded scans for each measurement timepoint for SOD1 animals.

| Timepoint [weeks] | 22-25 | 33 | 45 | 57-60 | 64 | 67 | 72-74 |
|---|---|---|---|---|---|---|---|
| $SOD1^{-/-}$ | 0% | 0% | / | 0% | 0% | / | 0% |
| Control | 0% | 0% | 14.2% | 0% | / | 0% | 12.5% |
